# SynNotch receptors for visualizing immunoreceptor force transmission and downstream signaling in vivo

**DOI:** 10.64898/2026.06.26.734558

**Authors:** Menglan Li, Jintian Lyu, Kaitao Li, Ameya Dravid, Deepali Balasubramani, Amir H. K. Ashkezari, Hyun-Kyu Choi, Gabriel A. Kwong, Ankur Singh, Cheng Zhu

## Abstract

Immunoreceptors experience forces that modulate their activities; however, demonstrating this in vivo has been limited by technical challenges. As a first step toward meeting this challenge, we adapted a synthetic Notch (SynNotch) receptor system to report force transmission through immunoreceptors in vivo by replacing the native ligand-binding domain with a receptor-specific antibody and rewiring Notch signaling to drive EGFP or luciferase expression. We expressed SynNotch on Jurkat cells targeting CD40 or T cell receptor (TCR) and characterized their activation in coculture with B or T cells, defining the requirements, optimal conditions, and kinetics of activation. Using complementary mechanobiology approaches, we quantified the exogenous force required for reporter activation and verified that activation depends on forces generated by receptor-expressing sender cells rather than SynNotch-expressing receiver cells. By implanting sensors and targets into immunocompromised mice, we visualized mechanically activated reporter expression on CD40 and TCR-targeting SynNotch cells in vivo. Furthermore, CD40 and TCR signaling was amplified when the receptor bore force against mechanical support from immobilized ligand, indicating that force functions as biologically relevant co-stimulus. Together, our results establish mechanically activated SynNotch reporters as a useful strategy for detecting receptor-associated mechanical signaling across 2D coculture, 3D organoid, and in vivo systems.

## INTRODUCTION

In cell biology, receptor signaling typically occurs when ligands, either soluble or immobilized, bind to their corresponding receptors. When ligands are immobilized on solid surfaces, the receptor–ligand bonds may experience mechanical forces exogenously, endogenously, or both. Exogenously, forces externally applied to a cell, such as hydrodynamic forces of flowing fluids over the cell that is attached to a stationary surface or another cell, must be balanced by adhesive forces mediated by receptor–ligand bonds, which are transmitted through the receptor structure into the cell and supported by the cytoskeleton^1,2^. Endogenously, cells generate and exert cytoskeleton- and motor-dependent forces on their receptor–ligand bonds to mediate cell adhesion, power cell locomotion, organize macromolecular assemblies (e.g., forming an immunological synapse), and deform the substrate beneath the cells^3–8^. A fundamental premise of mechanobiology is that cells respond to signals encoded in mechanical forces, which can be received, transmitted, and/or transduced through their surface receptors^3,7^.

In the past two decades, several techniques have been developed to measure the endogenous forces exerted through specific receptor–ligand bonds. Traction force microscopy (TFM) technique immobilizes ligands on an elastic substrate with micro-beads embedded as displacement markers. Lateral forces applied by the cell on the receptors bound to ligands can be determined from the measured deformation field of the substrate using the theory of elasticity^9,10^. Alternatively, the substrate is made into micro-pillar array detectors (mPADs) to allow the determination of lateral forces from the measured deflection of the mPADs using the theory of a cantilever beam^11,12^. Molecular tension probe (MTP) technique connects the ligand to a solid surface through a DNA hairpin with a fluorophore-quencher pair attached to the two ends, which would be opened when the force applied by the cell on the receptor–ligand bond exceeds the designed threshold force to unfold the hairpin, thereby dequenching the fluorophore to enable visualization of force^13–16^. Alternatively, the DNA hairpin is replaced by a spider silk protein peptide of calibrated elasticity, and a fluorescence donor-acceptor pair replaces the fluorophore-quencher pair to allow the force to be determined from the measured changes in the Förster resonance energy transfer (FRET) efficiency due to the stretch of the peptide^17,18^.

Except for a modified MTP that anchors to the membrane using a cholesterol tag to enable measurement of molecular tension at cell-cell junctions^19^, all other aforementioned techniques require special surface preparation, making it dissimilar to the membrane of another cell, raising the concern that the receptor forces so measured may not resemble forces under physiological conditions. Although these techniques have been successfully employed to indicate that cells exert endogenous forces on their surface receptors bound to immobilized ligands in vitro, they have not yet been demonstrated to do so in vivo, as none of these methods can be readily implemented within a living organism. Building upon recently published methods^20–22^ based on the synthetic Notch (SynNotch) receptor engineering^23^, we repurposed the technology platform of the Lim lab^23^ to develop a mechanically activated transcriptional reporter system for recording in vivo force on two specific immunoreceptors: CD40 on B cells and T cell receptor (TCR) on T cells. Notch is a cell surface receptor consisting of a negative regulatory region sandwiched by a ligand-binding domain and an intracellular domain, which can be activated by external force^24^. Activation involves three cleavages and nuclear translocation to upregulate the transcription of target genes^25^. The properties of Notch have been exploited to construct chimeric molecules in syntenic biology with various applications^26^. Similar to the published methods^20–22^, our adaptation of this platform is based on the mechano-sensitive and -responsive properties of the Notch receptor.

Our design uses the single-chain variable fragment (scFv) of an antibody against either human CD40 or mouse TCR to replace that of the anti-CD19 antibody of the published SynNotch construct^23^, but keeps the rewiring of the Notch signaling circuit to induce expression of a reporter—either an enhanced green fluorescence protein (EGFP) or a luciferase enzyme. Specific binding of SynNotch to target receptor induces sender cell to exert force on the receptor, which is transmitted to the SynNotch to activate the reporter expression in receiver cell, and the resulting accumulated transcriptional readout can be detected microscopically or by flow cytometry. Upon implanting SynNotch-expressing receiver cells in animals, allowing them to interact sender cells expressing the target receptor, and monitoring the resulting reporter expression, we visualized SynNotch activation induced by force transmitted through specific immunoreceptor in vivo. We exemplified the design, in vitro characterization, and in vivo validation of this mechanically activated transcriptional reporter using Jurkat cells as receiver cells, B cells or T cells as the sender cells, and CD40 or TCR as target immunoreceptors, illustrating the utility and adaptivity of the repurposed SynNotch system.

We constructed our specific SynNotch system to target two important immunoreceptors. CD40 is a key costimulatory receptor on B cells, and its binding to CD40 ligand (CD40L) delivers complementary signals alongside B cell antigen receptor (BCR) signaling to prevent B cell silencing or deletion^27^. Our recent in vitro studies showed that B cells exert tension on CD40–CD40L bonds, and force enhances CD40 signaling as well as antibody class-switch^16^. The TCR is arguably the most important receptor of T cell-mediated adaptive immunity because its interaction with peptide presented by the major histocompatibility complex (pMHC) molecule allows antigen recognition by the T cell, leading to its activation, proliferation, differentiation, and effector functions^28–30^. Several studies using DNA-based MTP have reported endogenous forces of 10-20pN on TCR engaged with pMHC or anti-TCR antibody^13,31–34^. Other studies using peptide-based MTP observed much lower forces on TCR^17,35^. Importantly, one of the major models of TCR triggering and antigen recognition is the mechanosensor model, which relies on the dual hypothesis that 1) TCR experiences mechanical forces in vivo and 2) such forces are biologically relevant co-stimuli that T cells detect and interpret to guide immune signaling^3,36^. The first hypothesis has been supported by the present in vivo experiments, finding SynNotch activation on receiver cells, respectively targeting CD40 on B cells and TCR on T cells in living animals. Moreover, we observed that such forces amplify activation of B cells and T cells mediated by CD40 and TCR, respectively, supporting the second hypothesis.

## RESULTS

### Construction of SynNotch with anti-hCD40 or anti-mTCR scFv

The gene circuit design of SynNotch has been used in multiple studies^20,23^. Our particular design is illustrated in Figure 1A (left). Initially, a response element was transduced into Jurkat T cells for dual purposes: firstly, to constitutively express a fluorescent protein (Blue Fluorescent Protein/mCherry), thereby enabling the distinction between receiver and sender cells; secondly, to reconstitute the GAL4-VP64 transactivation system to facilitate the expression of a reporter (or luciferase). Subsequently, the same Jurkat cells were transduced with an on-membrane construct comprising an anti-human CD40 (αhCD40) or anti-mouse TCR (αmTCR) scFv domain to replace the original Notch ligand-binding domain to recognize the target receptor and a Notch core negative regulatory region (NRR) that connects to an intracellular domain (ICD) (Figure 1A, middle). Similar to canonical Notch signaling^37^. Specific binding of the antibody scFv to the target receptor induces the sender cell to generate and exert force on the target receptor, which transmits to the bound SynNotch molecule to trigger a regulated intramembrane proteolysis (RIP) in the receiver cell. In this two-step process, a metalloprotease from the ADAM family cleaves SynNotch at the S2 site, and a γ-secretase cleaves the transmembrane domain (TMD), thereby releasing it to enter the nucleus, where it interacts with a response element to activate the expression of EGFP or Luciferase^38^ (Figure 1A, right). Thus, SynNotch activation in receiver cells provides a mechanically activated transcriptional readout of force transmission from sender cells through the target receptor.

**Figure 1.**
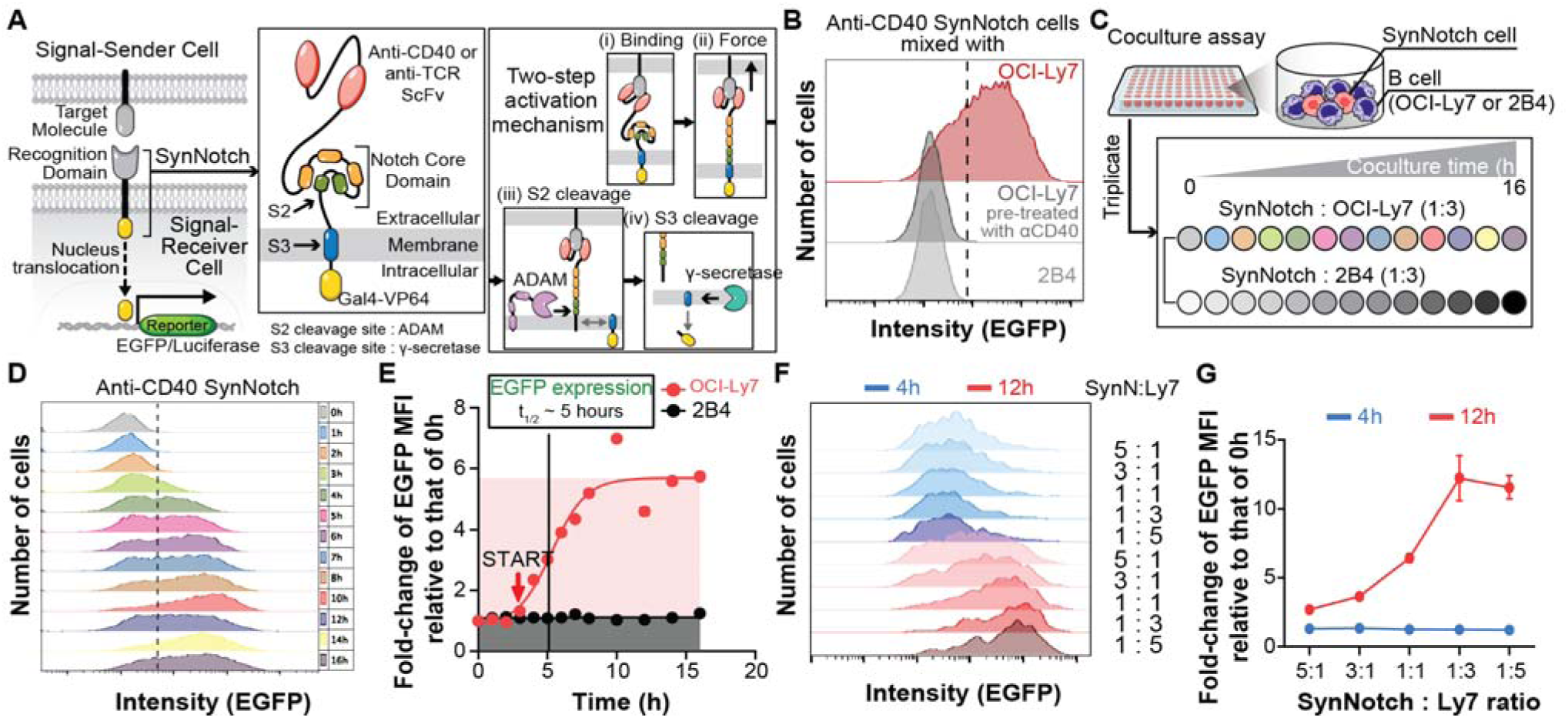
SynNotch configuration and αhCD40-SynNotch activation kinetics in a 2D coculture system. (A) *Left:* Schematics of detecting forces on CD40 or TCR using the SynNotch system. *Middle:* Zoom-in to our SynNotch system, which consists of a recognition domain for the target molecule, which is an antibody scFv against either human CD40 (αhCD40) or mouse TCR (αmTCR), a native notch core NRR that connects to a Gal4-VP64 intracellular domain. *Right:* Binding of αhCD40/αmTCR on sender cell (i) enables unfolding of NRR domain by force (ii), which expose S2 site by metalloproteases of the ADAM/TACE family (iii). S2-cleaved SynNotch then undergoes cleavage at S3 site by γ-secretase, allowing for the translocation of the intracellular domain into the nucleus to induce reporter EGFP expression^24^ (iv). The response element also constitutively expresses a blue fluorescent protein/mcherry to distinguish receiver cells from sender cells. (B) Histograms of EGFP fluorescence in receiver cells (Jurkat) expressing αhCD40-SynNotch cocultured with 3X excessive number of sender cells (B lymphoma cell line OCI-Ly7) expressing CD40 in the absence or presence of an anti-CD40 blocking antibody, or control cells (2B4) not expressing the target receptor. (C) 96-well plate layout and experimental procedure used to generate the results in (D) and (E). (D) αhCD40-SynNotch expressing Jurkat cells were cocultured with OCI-Ly7 or 2B4 cells at a 1:3 receiver-to-sender/control cell ratio in different wells at various times. Cells were harvested and their EGFP expressions were measured by flow cytometry and plotted as fluorescence histograms at the indicated times. (E) Mean ± SEM of Fold-change of EGFP mean fluorescence intensity (MFI) relative to that of 0 h (N = 3, n > 10,000 cells) at various times, measured from samples that were cocultured 5×10^4^ αhCD40-SynNotch expressing cells with 1.5×10^5^ OCI-Ly7 cells (green) or 1.5×10^5^ 2B4 cells (black). The data of the experimental group are fitted to a sigmoidal model. (F) Representative fluorescence histograms showing EGFP expressions in SynNotch-expressing cells activated by OCI-Ly7 cells at different mixing ratios. All 1.5×10^5^ cells were fixed across all conditions. Blue and red histograms in the upper and lower panels were measured 4- and 12-h post-coculture, respectively. (G) Mean ± SEM with individual data points (N = 3, n > 10,000 cells) of fold-change of EGFP MFI relative to that of the control group at different ratios for 4-h (blue) and 12-h (red) post-coculture. Note that the individual data points are obscured due to overlapping.

The specificity of SynNotch activation was first tested by mixing the receiver cells (Jurkat) with sender cells (human B lymphoma cell line OCI-Ly7 or primary mouse CD8^+^ T cells) that express the target receptor (CD40 or TCR, depending on the specific SynNotch scFv) in the absence or presence of an antibody to block the SynNotch binding, or with control cells not expressing the target receptor (mouse T hybridoma cell line 2B4 or OCI-Ly7).

After 8-h, significant EGFP expression was observed exclusively in SynNotch-expressing receiver cells cocultured with target receptor-expressing sender cells, but not with control cells lacking the target receptor (Figure 1B and Figure S1A). Furthermore, treatment with an anti-CD40 blocking antibody abolished EGFP expression in αhCD40-SynNotch-expressing Jurkat cells cocultured with CD40-expressing OCI-Ly7 cells (Figure 1B). These data confirm that reporter expression was upregulated through specific interactions between the designated SynNotch antibody scFv and the target receptor.

### Reporter activation kinetics and sensitivity of SynNotch in a 2D coculture system

To characterize the reporter activation kinetics of SynNotch, receiver cells and sender cells were sequentially combined at staggered time points in ratios of 1:3 (for the αhCD40-SynNotch) or 1:5 (for the αmTCR-SynNotch). The cell mixtures were subsequently centrifuged to facilitate cell-cell interaction within wells of a 96-well plate, thus establishing a two-dimensional (2D) coculture system (Figure 1C). All samples were collected simultaneously at the 16-hour mark, corresponding to the predefined incubation durations for each coculture condition. Subsequently, cells were analyzed via flow cytometry to assess cell viability and EGFP expression. No significant Jurkat cell death was observed up to 16 h of coculture (Figure S2). Fluorescence histograms (Figure 1D and Figure S1B) were quantified by their mean fluorescence intensities (MFI). The data at later time points were normalized to the initial MFI, plotted against time, and the trend was modeled using a sigmoidal curve to illustrate activation kinetics (Figure 1E). EGFP expression became detectable around 3 h after coculture, reached near-maximal levels at approximately 10 h, and then plateaued, with a half-maximum time (*t*_1/2_) of about 5 h (Figure 1E and Figure S1C). The specificity of αhCD40-SynNotch activation was confirmed by the absence of target-independent activation when Jurkat cells were cocultured with control sender cells (2B4) not expressing CD40 at all tested time points (Figure 1E).

We further tested the sensitivity of our reporter system by measuring SynNotch activation with a range of mixing ratios of receiver cells to sender cells. No significant difference in the αhCD40-SynNotch activation levels was observed across the different ratios tested at 4-h post-mixing coculture (Figures 1F and 1G), which represents the onset of activation at early time points (Figures 1D and 1E). However, after 12-h coculture, increasing the proportion of sender cells relative to receiver cells gradually elevated EGFP expression, reaching a plateau when the sender cells were three times more than the receiver cells (receiver/sender ratio of 1:3) (Figures 1F and 1G). In comparison, αmTCR-SynNotch activation peaked at a receiver/sender ratio of 1:5 (Figure S1D).

### Response of αhCD40-SynNotch reporter activation to calibrated externally applied forces

To more precisely calibrate SynNotch reporter activation under defined external forces than previous reports^25,39,40^, we employed our newly developed parallel magnetic force activation (PMFA) assay^16^ to apply a range of calibrated forces on αhCD40-SynNotch, and measured the corresponding reporter expression. The PMFA assay is a high-throughput platform^16^ adapted from a previously published system^41^. The device consists of a 24-well plate, the lid of which includes a 3D-printed housing to mount two magnets in an antiparallel configuration with a calibrated distance above the bottom surface (Figure 2A). Jurkat cells expressing αhCD40-SynNotch were seeded at the bottom of each well, with a 5 mm diameter coverslip coated with poly-L-lysine (PLL) affixed to the center. Paramagnetic beads coated with varying densities of soluble hCD40 were added to facilitate binding to the cell surface αhCD40-SynNotch. Closing the lid permitted the magnetic gradient to exert an average force of 25pN on each bead^16^, which was distributed over the individual hCD40–αhCD40-SynNotch bonds on the bead–cell interface. We used a biomembrane force probe (BFP) (Figure 2B)^16^ to measure the effective 2D affinity (Figure 2C, *A*_c_*K*_a_ where *A*_c_ is the contact area and *K*_a_ is the 2D affinity, both in μm^2^) and force-dependent bond lifetime (Figure 2D) of the hCD40–αhCD40-SynNotch interaction. Based on the αhCD40-SynNotch expression level (*m*_l_), we varied the coating density (*m*_r_) of the hCD40 on the beads (Figure 2E and Figure S3A) to achieve a range of values for the average number of bonds, calculated as a product of *m*_r_, *m*_l_, and *A K* ^42^. These are then translated into a range of forces per hCD40–αhCD40-SynNotch bond (Figure 2F and Figure S3B).

**Figure 2.**
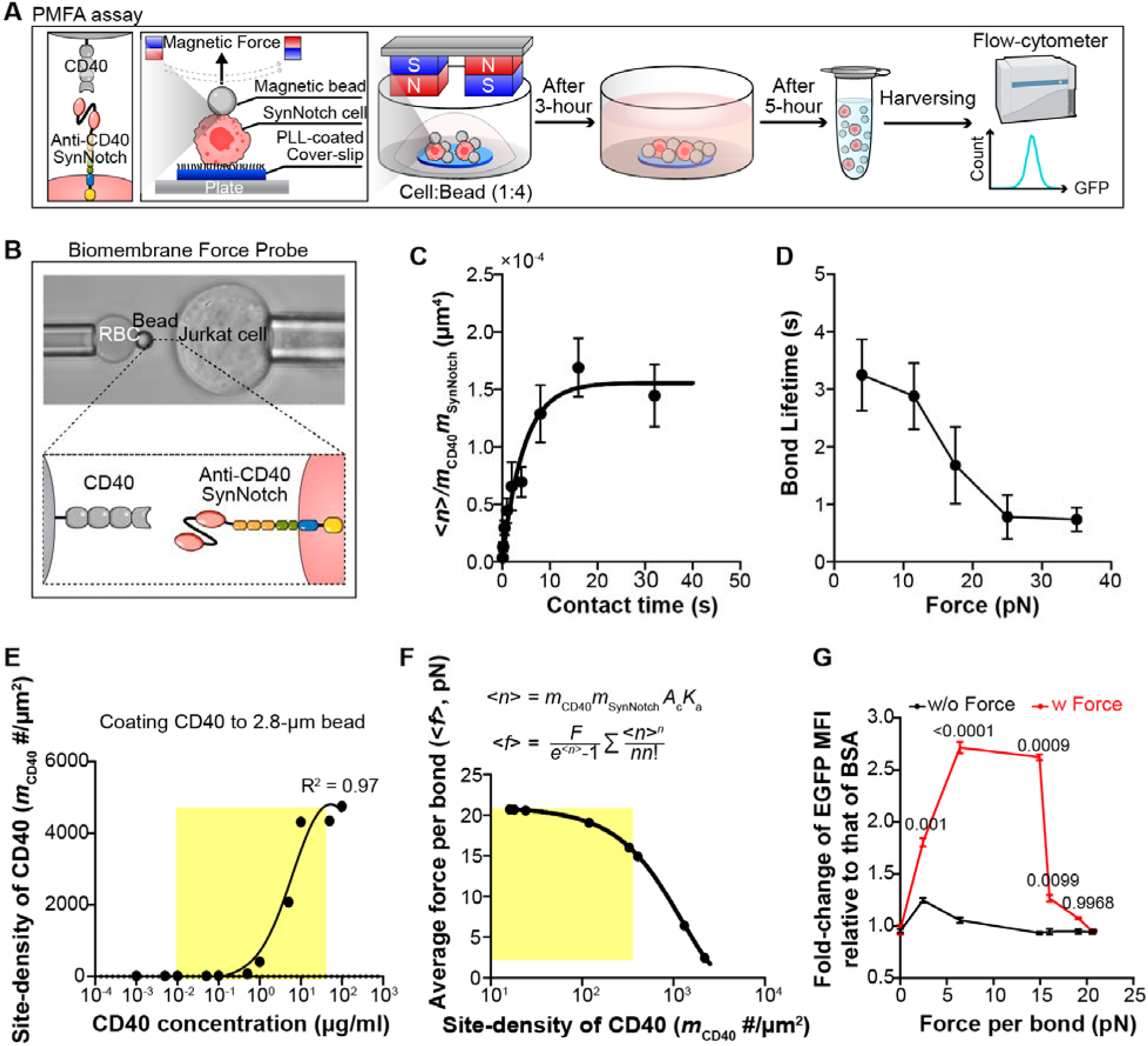
Response of αhCD40-SynNotch reporter activation to calibrated externally applied forces. (A) Working principle and procedure of the parallel magnetic force activation (PMFA) assay. SynNotch expressing Jurkat T cells were placed on PLL-coated circular coverslip (5-mm in diameter) at the center of each well in a 24-well plate onto which 4X excessive number of paramagnetic beads (2.8-μm in diameter) functionalized with soluble human CD40 protein were added. Cells were allowed to interact with beads via CD40–αhCD40-SynNotch bonds for 3-h in the presence (experimental) or absence (control) of an anti-parallel pair of 5-mm cubic neodymium magnets mounted on the plate lid after which the magnets were removed and more medium was added for an additional 5-h incubation. Cells were then collected and their EGFP expression levels were analyzed by flow-cytometry. (B) Photomicrograph (upper) and zoom-in view of the boxed area (lower) of the BFP setup. A biotinylated RBC is aspirated by a micropipette and a CD40-functionalized glass-bead is attached to its apex via biotin-streptavidin coupling (left). The RBC-bead assembly also acts as an ultrasensitive force sensor to detect binding of CD40 with αhCD40-SynNotch expressed on a Jurkat T cell aspirated by an opposite micropipette (right). (C) Mean ± SEM (n > 2 per point) of average number of bonds per contact «n)) normalized by site densities of CD40 (m_r_, in jxm”^2^) and ahCD40-SynNotch *(m^* in jxm’^2^) *vs* contact time (t_c_, in s). *(n) = —* ln(l — P_a_) where the adhesion frequency *P_a_* (= # binding events divided by total # of contacts) is directly measured from repeated bead-cell contacts. Data were fitted (curve) by a previously reported model^42^. The steady-state value is the effective 2D affinity (in jxm^4^). (D) Bond lifetime (mean ± SEM) *vs* force (n ≥ 15 per force bin) of CD40-coated beads interacting with Jurkat T cells expressing αhCD40-SynNotch. (E) Measurements (point, by flow cytometry) and trendline (curve) of CD40 site density *vs* CD40 coating concentration. The appropriate CD40 densities for the PMFA assay are highlighted. (F) Average force per bond (<*f*>) *vs* CD40 density (*m*_r_) determined using the indicated equations. In the upper equation, multiplying the CD40 density *m_r_* from the *x*-axis to the site density of αhCD40-SynNotch *m_l_* and the effective 2D affinity *A*_c_*K*_a_ measured in (C) yields the average number of bonds per contact <*n*>. The average force per bond *<f<* is then calculated using the lower equation from *<n>* and the average force per bead *(F)* applied the magnets in the PMFA assay. The appropriate *m_r_* values for PMFA assay are highlighted. Data (points) are fitted by a trend line (curve). (G) Fold change of EGFP MFI in Jurkat cells expressing αhCD40-SynNotch in contact with beads coated with hCD40 normalized by that of the same cells in contact with beads coated with BSA in the presence of soluble CD40 *vs* average force per bond for both experimental (red, with force) and control (black, without force) conditions. P-values (numbers) were determined using student t-test or one-way ANOVA, which gave the same results.

Utilizing the PMFA platform, we conducted a comparative analysis of αhCD40-SynNotch activation across two cellular groups, testing in both the absence and presence of a specified range of exogenous forces. The two groups maintained identical cell and bead preparations; however, the group with force employed a lid mounted with magnets, whereas the group without force utilized a standard 24-well plate lid devoid of magnets. Additionally, wells containing αhCD40-SynNotch-expressing Jurkat cells and BSA-coated beads with soluble hCD40 in the media served as another negative control for normalization in Figure 2G. The experiment began with a 3-h incubation during which the beads and cells were brought into contact to allow thorough interaction between hCD40 and αhCD40-SynNotch in the presence or absence of force. Subsequently, both lids with and without magnets were replaced by standard lids, and the wells were supplemented with additional media to facilitate EGFP induction reaching a steady expression. At the endpoint, cells were harvested and their EGFP expression levels were analyzed via flow cytometry. Under these conditions, αhCD40-SynNotch activation can be detected at forces as low as 2.4 pN per bond and reached maximal levels in the range of 6.4 to 15pN (Figure 2G). This calibration defines the range of externally applied forces that produce SynNotch activation after a defined stimulation period. Thus, the measured EGFP signal represents accumulated reporter output under the given force-loading conditions. To apply forces exceeding 15pN without further lowering the magnets—an adjustment complicated by the presence of media—it would be necessary to decrease the density of hCD40 on the paramagnetic bead. This reduction would decrease the average number of bonds with αhCD40-SynNotch on the cell to below one (Figure S3B), rendering them susceptible to rupture under force (Figure 2D) as beads were attracted to accumulate at the magnets, thus abolishing αhCD40-SynNotch activation (Figure 2G).

### Sender cell generates endogenous forces on SynNotch instead of the receiver cell

Generally, sender cells provide the force needed to activate Notch receptors during the endocytic process when Notch ligands engage with Notch receptors on receiver cells^25^. To test if Jurkat cells themselves exert endogenous forces on the transduced SynNotch and/or are capable of being activated by simply engaging with the target receptor without force acting on the SynNotch–receptor bond, we utilized a MTP for imaging piconewton forces^16,43^ (Figure 3A). The MTP comprises three single-stranded DNAs (ssDNA). The central region of strand 1 adopts a hairpin structure in the absence of force, with GC content carefully designed to regulate the threshold force required for unfolding. Its two termini serve as sequences for hybridization with complementary strands 2 and 3. One end of strands 2 and 3 is conjugated with a black hole quencher 2 (BHQ2) or a Cy3b fluorophore, respectively, such that after hybridization with strand 1 at the folded hairpin configuration the BHQ2 is adjacent to the Cy3b to quench its fluorescence. The other end of strand 2 is linked to a gold nanoparticle coated on cover glass, which further quenches the fluorophore. The other end of strand 3 is linked to a streptavidin, facilitating conjugation with a C-terminally biotinylated protein of interest such as soluble hCD40 or mTCR (Figure 3A, left), or control protein (αCD3 antibody as a positive control and BSA as a negative control). Receiver cells are placed on the MTP surface to facilitate SynNotch binding to target receptors and to enable endogenous force application to the molecular bonds. When the applied forces exceed the predetermined threshold, the hairpin unfolds, thereby separating the Cy3B from the BHQ2 and the gold nanoparticle, leading to the de-quenching of fluorescence (Figure 3A, right). To capture short-lived fluorescence signals terminated by bond dissociation and/or intermittent pulling by the cell, a single DNA strand complementary to the unfolded DNA hairpin (locker strand) is included in the media, which would hybridize with the hairpin upon its unfolding to lock it in the unfolded configuration, thereby accumulating the transient and discontinuous force signals over time^15^ (Figure 3A).

**Figure 3.**
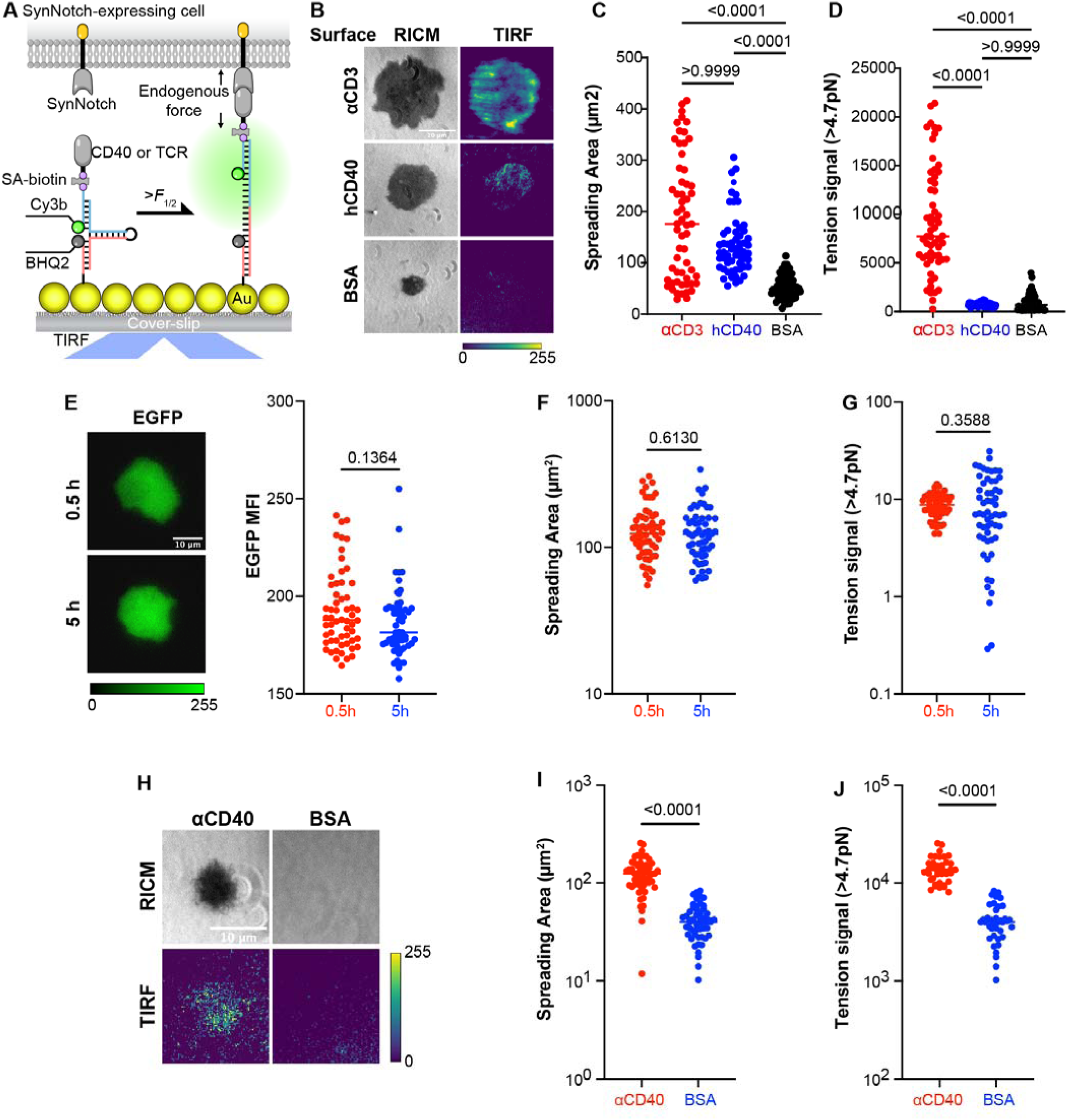
Force on αhCD40-SynNotch was generated by the sender cell instead of the receiver cell. (A) Schematic of MTP^43^. An MTP consists of a DNA hairpin to define a threshold force that has a 50% of probability to unfold it (*F*_1/2_), a fluorophore (Cy3b) strand and a quencher (BHQ2) strand. One end is linked to a gold nanoparticle-coated glass surface, and the other end is linked to a soluble hCD40, mTCR, anti-CD40 antibody, or anti-CD3 for positive control, or BSA for negative control. The Cy3b and BHQ2 are in close proximity when the hairpin is folded, thus the fluorophore is quenched. The hairpin unfolds when the cell exerts force greater than the threshold, separating BQH2 from Cy3b to generate a fluorescent signal. Adding a complementary ssDNA (locker) to hybridize with the opened hairpin prevents it from closing after bond dissociation and force removal, which enables the fluorescent signal to accumulate^43^. (B) Representative images by RICM and TIRF microscopy showing contact area and force signal of αhCD40-SynNotch expressing Jurkat cells 20 min after landing on coverslip surface functionalized by MTP tagged with indicated proteins. (C and D) Quantification of spreading area (C) and tension signal (D) exemplified in (B). Each point represents a cell, n = 55, 58, 48. (E) *Left*: Representative EGFP fluorescent images of an αhCD40-SynNotch expressing Jurkat cell at the indicated time post-cell seeding. *Right:* Quantification of EGFP MFI for conditions on the left. Each point represents a cell, n = 58, 55. (F and G) Quantification of spreading area (F) and tension signal (G) in (E). Each point represents a cell, n = 58, 55. (H) Representative images by RICM and TIRF microscopy of OCI-Ly7 cells 20 min after landing on coverslip surfaces functionalized by MTP tagged with indicated proteins. (I and J) Quantification of spreading area (I) and tension signal (J) exemplified in (H). Each point represents a cell, n = 55, 51. All MTP experiments were repeated three times. P-values (numbers) were determined using student t-test (for two conditions) or one-way ANOVA (for three conditions).

We used reflection interference contrast microscopy (RICM) and total internal reflection fluorescence (TIRF) microscopy to visualize, respectively, cell spreading and pulling on the MTP surfaces (Figure 3B, Figure S4A). Quantification of the RICM images shows that SynNotch-expressing Jurkat cells spread significantly smaller areas on the negative control surfaces functionalized with MTP tagged with the irrelevant protein BSA than on the experimental surfaces functionalized with MTP tagged with the target receptors (hCD40 or mTCR), which were comparable to spreading on the positive control surfaces functionalized with MTP tagged with αCD3 (Figure 3C and Figure S4B). These results indicate that the significantly larger areas of Jurkat cell spreading on the target receptors (hCD40 and mTCR) were mediated by their specific interactions with the αhCD40-SynNotch or αmTCR-SynNotch on the receiver cells, just like the Jurkat cell spreading on the αCD3 was mediated by its specific interaction with the TCR-CD3 constitutively expressed on Jurkat T cells.

Interestingly, interactions of SynNotch-expressing Jurkat cells with their respective target receptors (hCD40 or mTCR) coated on MTP surfaces did not generate endogenous forces during the approximately 30-minute observation period. This was evidenced by the lack of significant difference in the Cy3b fluorescence levels between the experimental surfaces and the negative control surfaces coated with MTP tagged with the irrelevant protein BSA (Figure 3D and Figure S4C). Importantly, Cy3b signals from the surfaces functionalized with MTP tagged with αCD3 were substantially higher (Figure 3D and Figure S4C), indicating that specific interactions of Jurkat T cells with MTP surfaces induced endogenous forces to bear on TCR-CD3–αCD3 bonds. This positive control, plus our recent report that Jurkat cells stimulated to express CD40L pull on CD40-tagged MTP^16^, rule out the possibility that the Jurkat cells might be unable to generate forces, and instead indicate the inability of force induction specific to SynNotch.

### Target receptor on the acellular surface could not induce detectable SynNotch activation in a short period

Binding of soluble ligands to the cell surface Notch does not activate Notch signaling because ligand binding in solution does not generate the necessary force on Notch^25^. However, cell movement may exert force on cell surface receptors that are engaged with immobilized ligands^3^. To assess whether SynNotch can be activated through engagement of target receptors immobilized on plastic surfaces, we measured the change in fluorescent intensity in Jurkat cells positioned on hCD40-tagged MTP surfaces over a period of 0.5 to 5 h. This measurement was based on our earlier observation that EGFP upregulation begins approximately at 3 h and reaches half of its maximum level by around 5 h post-coculture (Figure 1E). Image quantification revealed statistically indistinguishable EGFP fluorescent intensities (Figure 3E), spreading areas (Figure 3F), and tension signals (Figure 3G) of Jurkat cells on hCD40-tagged MTP surface between 0.5 and 5 h, finding no detectable SynNotch activation over the 5-h period, likely due to the lack of receiver cell pulling despite the continuous interaction with αhCD40. This data suggests that, within a short period of 5 h, receptor binding to αhCD40-SynNotch upregulates EGFP reporter expression when hCD40 is expressed on live sender cells, but not when it is coated on acellular surfaces. This is likely due to the ability of live cells to generate and exert forces on the αhCD40-SynNotch–hCD40 bonds, a capability that inert plastic surfaces lack. However, after αmTCR-SynNotch expressing Jurkat cells were placed on MTP surface tagged with soluble mTCR for 14 h, flow cytometric analysis of the harvested cells revealed SynNotch activation much higher than the negative control (the same Jurkat cells cocultured with TCR-free Ramos cells) but at similar levels as the positive control (the same Jurkat cells cocultured with mouse CD8^+^ T cells) (Figure S4D). The observed SynNotch activation on the acellular surfaces is consistent with the recent report that microparticle-conjugated and surface microcontact-printed target receptors activate SynNotch over 24 hours or longer^44^, and might result from force on the αmTCR-SynNotch– mTCR bonds due to migration of Jurkat cells on ECM proteins that they secreted over the prolonged period of time.

Previous works by us and others have shown that B and T cells exert endogenous forces on CD40L^16^ and TCR^13,15,31,32^, respectively. To verify that OCI-Ly7 cells also exert endogenous forces on αhCD40, they were placed on surfaces functionalized with αhCD40-tagged MTP and imaged by RICM and TIRF. The results revealed a significantly increased spreading area and tension signals compared to cells positioned on control surfaces coated with BSA-tagged MTP, as illustrated in Figures 3H-3J. This confirms that the interaction between CD40 and αhCD40 indeed induced OCI-Ly7 cells to generate endogenous forces to bear on the CD40–αhCD40 bonds.

### Reporter activation kinetics of SynNotch in an organoid system

As an intermediate step toward in vivo testing, we utilized an organoid system to coculture SynNotch expressing receiver cells and target receptor expressing sender cells in 3D, which better mimics their interactions under physiological conditions compared to the previous 2D coculture system (Figure 1). The engineered 3D organotypic lymphoid culture system, initially developed to study human germinal center reactions in B cells^45^ and human lymphoma growth^46^, serves as an in vitro immune organ module capable of customization to mimic in vivo environments. Using these lymphoid organoids, we have previously demonstrated that CD40L–CD40 ligand–receptor interactions amplify the B cell receptor (BCR)–MYD88–TLR9 multiprotein super-complex and induce cooperative signaling pathways in lymphoma cells, which reduce the efficacy of compounds targeting the BCR pathway members^46^. Receiver (Jurkat) cells expressing αhCD40-SynNotch and sender (OCI-Ly7) cells expressing CD40 or control (2B4) cells not expressing the target receptor were mixed at 1:3 ratio in complete cell culture medium plus maleimide functionalized polyethylene glycol (PEG-4-MAL) and loaded into wells of 96 well (Figure 4A, left), which underwent gelation (Figure 4A, middle) in 15 min to entrap the cell mixture into a 3D organoid per well (Figure 4A, right). As in the previous 2D experiment (Figure 1B), we confirmed specificity of αhCD40-SynNotch activation in the 3D system: Jurkat cells expressed EGFP when cocultured with OCI-Ly7 but not with 2B4 cells, and this EGFP expression was eliminated by an anti-CD40 blocking antibody in receiver cells cocultured with sender cells (Figure 4B). Similarly, specific αmTCR-SynNotch activation in the 3D system was demonstrated as EGFP expression was significantly higher in receiver cells cocultured with T cells compared to TCR-free Ramos cells (Figures S5A-S5B). The αhCD40-SynNotch activation in the 3D system followed a slower kinetics than the 2D system (Figure 4C). Although detected as early as 3 h, EGFP expression reached near-maximal levels at 24 h, with a reporter half-maximum time *t*_½_ ≈ 13 h (Figure 4D), more than doubling the 5-h half time of the 2D system (Figure 1E). This slower reporter activation may reflect several non-mutually exclusive features of the organoid environment. Compared with the 2D coculture system, sender and receiver cells in the hydrogel may require more time to migrate, form productive contacts, and interact with one another through receptor–SynNotch bonds. In addition, reduced nutrient or oxygen diffusion, lower effective local cell density, altered cell distribution within the matrix, and matrix stiffness-dependent effects on cell migration or signaling may also contribute to the delayed reporter activation. Additionally, we noted that in the 3D system, the EGFP expression remained at the peak level for another 24 hours with a mild decline thereafter. This prolonged activity may also be attributed to the inherent characteristics of the organoid system, which potentially replicate physiological conditions more accurately than the 2D system, thereby enhancing the maintenance of cellular function over an extended period.

**Figure 4.**
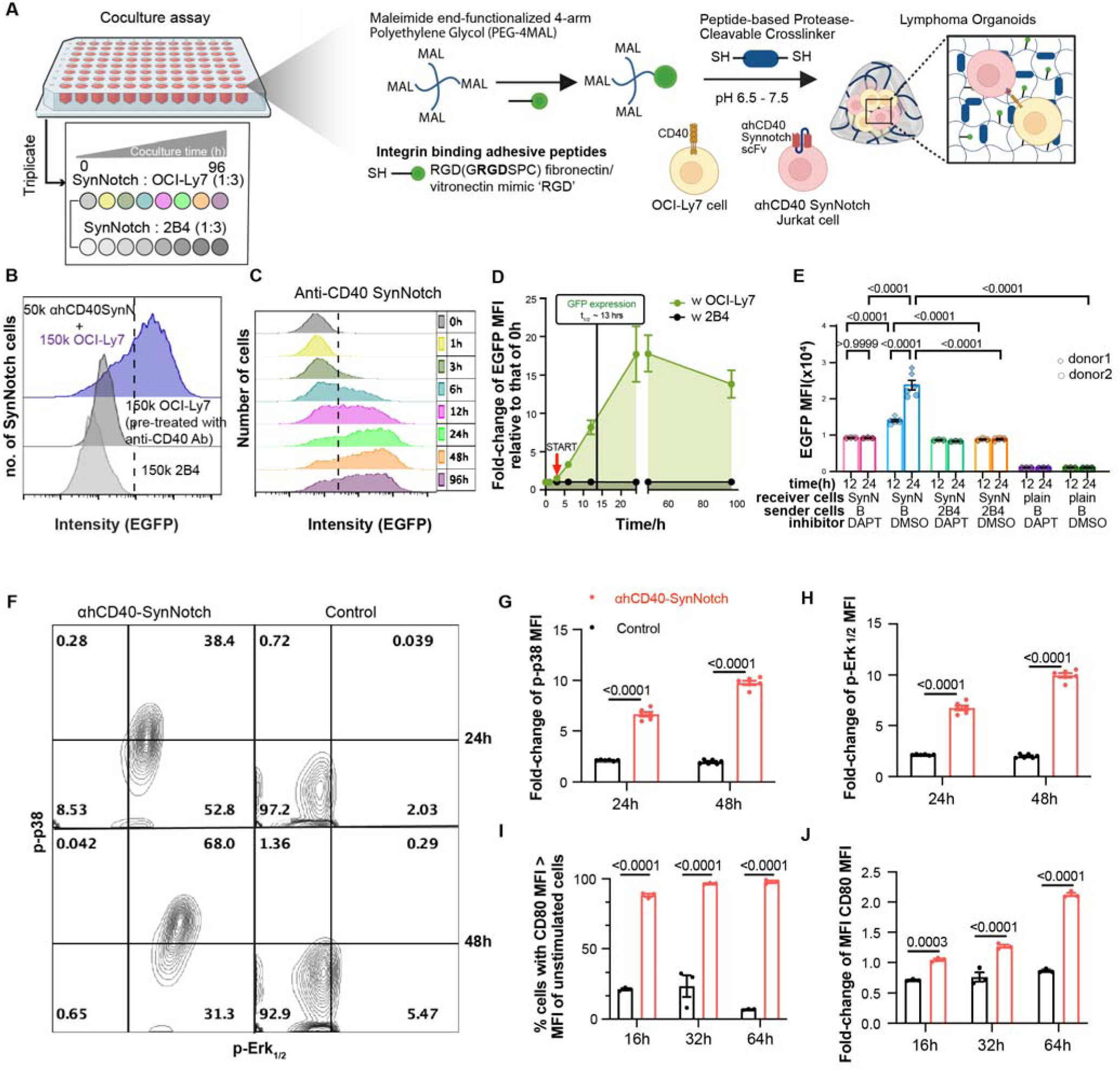
Reporter activation kinetics of αhCD40-SynNotch in a 3D organoid coculture system. (A) *Left*: 96-well plate layout and experimental procedure used to generate the results in (C) to (I). αhCD40-SynNotch expressing Jurkat cells and CD40 expressing OCI-Ly7 cells or 2B4 not expressing CD40 (negative control) were mixed in a 1:3 ratio and loaded onto wells of a 96-well plate at different times to achieve the indicated coculture times in organoids. At the endpoint, organoids were degraded, cells were harvested, stained with appropriate antibodies, and analyzed by flow cytometry for EGFP expression (in receiver cells) as well as CD80, p-p38 and p-Erk1/2 upregulation (in sender cells). *Middle:* Illustration of how organoids are made. *Right*: Schematic of an organoid and the encapsulated cells. (B) Representative fluorescence histograms of the specificity control experiment, showing right-shift of EGFP fluorescence in receiver cells cocultured with sender cells in the absence (blue) relative to that in the presence (black) of blocking antibody, or relative to that cocultured with control cells (gray). (C) Representative fluorescence histogram stack of EGFP upregulation in αhCD40-SynNotch expressing Jurkat cells at various times. The EGFP+ gate was defined using the same receiver cells without interacting with the sender cells. (D) Mean ± SEM (N=5, n>10,000) of EGFP MFI of receiver cells cocultured with sender (green) or control (black) cells at various times. Red arrow with START indicates the starting point of EGFP upregulation. (E) αhCD40-SynNotch Jurkat cells or plain Jurkat cells were pretreated with 10 μM DAPT or DMSO for 4 h or and then cocultured with primary human B cells or 2B4 cells in a ∼1:3 ratio. Cells were collected after 12 and 24 h and analyzed by flow cytometry for EGFP expression. Mean ± SEM with individual measurements show two independent experiments using B cells isolated from two human donors. P-values were calculated using one-way ANOVA. (F) Representative scatterplots of p-p38 vs p-Erk1/2 fluorescence staining of sender (OCI-ly7) cells cocultured for 24 h (upper row) or 48 h (lower row) with receiver (αhCD40-SynNotch expressing Jurkat) cells (left column) or control (plain Jurkat) cells (right column). (G and H) Mean ± SEM with individual measurements of fold-change of p-p38 (G) or p-Erk1/2 (H) MFI in sender cells cocultured for 24 or 48 h with receiver cells (red) or control (black) cells relative to those of control cell. (I and J) Mean ± SEM with individual measurements of fold-change of CD80^+^ cell frequency (I) and MFI (J) in sender cells cocultured for 16, 32 or 64 h with receiver (red) or control (black) cells relative to those of control cells. P-values in (E) and (G) to (J) were calculated using one-way ANOVA.

To further validate αhCD40-SynNotch activation in a more physiologically relevant model than the Large B Cell Lymphoma (DLBCL) line OCI-Ly7 harboring MYD88 and CD79B mutations that constitutively activate NF-κB, and to test the dependence of reporter expression on the SynNotch TMD cleavage by γ-secretase, we performed additional 3D organoid experiments using primary human B cells isolated from peripheral blood of two healthy donors. CD19-positive selected B cells were encapsulated in the 3D organoid with αhCD40-SynNotch Jurkat cells pretreated with 10μM N-[N-(3,5-Difluorophenacetyl)-L-alanyl]-S-phenylglycine t-butyl ester (DAPT) for four hours at 37 °C to inhibit γ-secretase activity, and DAPT was maintained throughout the coculture for 12 or 24 h.

Primary B cells induced robust EGFP upregulation in DMSO-treated receiver cells, and reporter expression increased from 12 to 24 h (Figures 4E). In contrast, DAPT treatment strongly reduced SynNotch activation in receiver cells. Importantly, the EGFP expression levels on the DAPT-treated receiver cells cocultures with CD40-expressing B cells and 2B4 cells not expressing CD40 were statistically indistinguishable. Regardless of the DAPT or DMSO treatment, no EGFP expression was detected on plain Jurkat cells not expressing SynNotch cocultured with B cells. Together, these data extend the OCI-Ly7 coculture findings to primary human B cells and confirm that mechanically activated αhCD40-SynNotch reporter expression requires γ-secretase-mediated TMD cleavage.

### SynNotch engagement activates sender cells in vitro

As a key interaction mediating intercellular interaction between T and B cells, CD40–CD40L binding is known to activate B cell signaling, promote sustained proliferation, expansion, differentiation, and antibody isotype switching^27^. We therefore asked whether B cells, which are the sender cells in our experiment, were also activated by interaction of their CD40 with the αhCD40-SynNotch on the receiver Jurkat T cells. Following our previous in vitro studies using p38 and Erk1/2 phosphorylation as B cell activation markers^16^, we examined the phosphorylation of these critical molecules in the B cell signaling pathways. OCI-Ly7 cells expressing CD40 were cocultured in organoids with Jurkat cells that either expressed or did not express (control) αhCD40-SynNotch. OCI-Ly7 cells were collected at various time points, permeabilized, stained with antibodies against phosphorylated p38 (p-p38) and Erk1/2 (p-Erk1/2), and analyzed via flow cytometry. The fluorescence scattergrams (Figure 4F) and mean fluorescence intensity (Figures 4G and 4H) showed a significant increase in phosphorylation levels of p38 and Erk1/2 at both 24 and 48 h in OCI-Ly7 cells cocultured with Jurkat cells expressing αhCD40-SynNotch. In contrast, no such increase was observed in OCI-Ly7 cells cocultured with control Jurkat cells lacking αhCD40-SynNotch or CD40L. These results suggest that, similar to CD40L, engagement of αhCD40-SynNotch with B cell CD40 triggers B cell signaling. Furthermore, we analyzed the expression of CD80, a surface marker for B cell activation and ligand for the key co-stimulatory molecule CD28 on T cells. Consistent with the upregulation of p-p38 and p-Erk1/2, CD80 expression was significantly upregulated in OCI-Ly7 cells in the experimental group cocultured with Jurkat cells expressing αhCD40-SynNotch, but not in the control group cocultured with plain Jurkat cells (Figures 4I-4J).

Another key molecular interaction between T and B cells is the TCR–pMHC binding, which triggers T cell signaling, effector functions, and more differentiation^3^. Therefore, we hypothesized that primary CD8^+^ T cells cocultured with Jurkat cells expressing αmTCR-SynNotch in 3D organoids would induce upregulation of T cell activation markers. Supporting our hypothesis, flow cytometric analysis revealed significantly increased expression CD44 and PD-1 on mouse T cells after 48 h of 3D coculture with αmTCR-SynNotch expressing Jurkat cells compared to the control group that used plain Jurkat cells (Figures S5C and S5D). Together, these results indicate that not only does SynNotch engagement with CD40 and TCR induce respective B and T cells to generate and exert forces on the resulting bonds to activate reporter expression on the Jurkat cells, but these forces also induce CR40 and TCR-mediated activation of B and T cells, indicating the biological relevance of these forces.

### Permitting endogenous force on CD40–CD40L bonds enhances B cell activation

In the natural process of Notch activation in receiver cells, sender cells pull on Notch ligand via an endocytic process upon its engagement with the Notch receptor^24,47,48^. Activated CD4^+^ T cells transfer CD40L to B cells through interaction with CD40, which also occurs via endocytosis^49^. We have recently shown that T cells exerted endogenous force on CD40L, applying exogenous force on CD40 amplifies B cell activation, and limiting force on CD40–CD40L bonds suppresses B cell signaling^16^. To further validate the role of mechanical force on CD40 in B cell activation, we utilized tension gauge tethers (TGTs)^16,50^ tagged with CD40L to compare CD40-induced B cell activation when the endogenous forces on the CD40–CD40L bond were permitted to be as high as 56pN with that when such forces were limited to be no more than 12pN. TGT is similar to MTP in that both take advantage of the property of double-stranded DNA (dsDNA) oligos to dissociate above a tunable force threshold (compare Figure 3A and Figure 5A, right). However, MTP utilizes a ssDNA that forms a hairpin with a closed end, which unfolds under forces above a threshold and can refold reversibly after force removal (unless refolding is prevented by hybridization with a locker strand) (see Figure 3A). In contrast, TGT employs a dsDNA duplex with open ends (Figure 5A, right), which breaks irreversibly when force exceeds the threshold, making re-connection unlikely even after force is removed. TGT rupture releases CD40s from experiencing forces above the threshold and prevents them from binding CD40L tagged on remaining TGTs. This reduces the number of free CD40 on the cell surface and the available TGT-tagged CD40L for binding. Consequently, the TGT assay enables us to study how endogenous forces influence B cell activation mediated by CD40–CD40L bonds.

**Figure 5.**
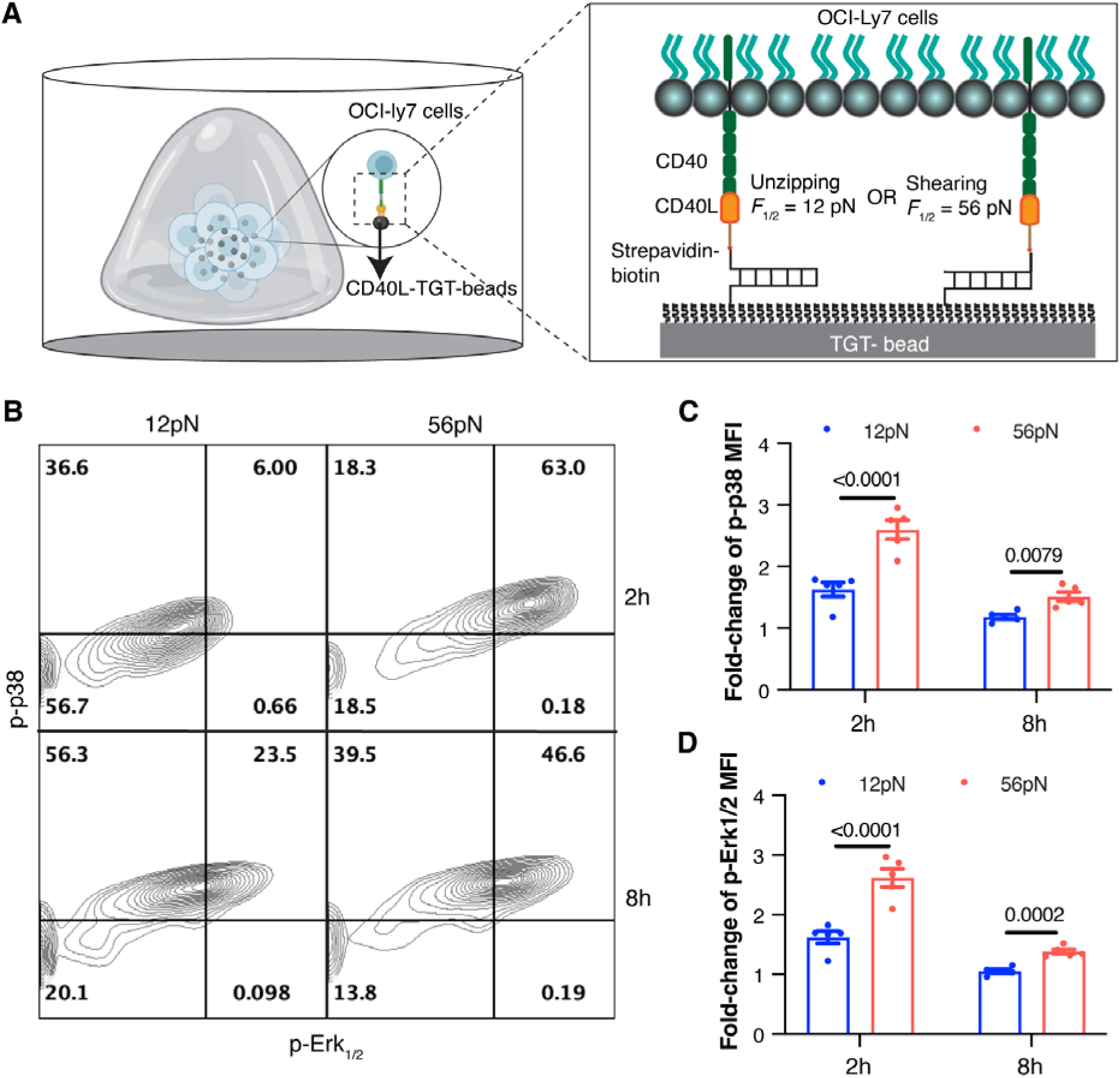
Permitting endogenous force on CD40 enhances B cell activation. (A) *Left*: Schematic of an organoid and zoom-in view of encapsulated OCI-ly7 cells and beads functionalized with TGT-tagged CD40L. *Right:* Zoomed-in view of the box on the left showing a schematic of TGT and its working principle. TGTs with two different threshold forces were depicted, depending on the mode of dsDNA separation: Force to achieve 50% chance of separation (*F*_½_) is 12pN for the unzipping mode (*left*) and 56pN for the shearing model (*right*). The overall experiment procedure is similar to Fig. 4 except that the receiver cells were replaced by beads coated with TGT-tagged CD40L (experimental) or BSA (control). (B) Representative scatterplots of p-p38 vs p-Erk1/2 fluorescence staining of OCI-ly7 cells cocultured for 2 h (upper row) or 8 h (lower row) with CD40L TGT beads of 12pN (left column) or 56pN (right column) threshold force. (C and D) Mean ± SEM with individual measurements of fold-change of p-p38 (C) or p-Erk1/2 (D) MFI in OCI-Ly7 cells cocultured for 2 or 8 h with CD40L TGT beads of 12pN (blue) or 56pN (red) threshold relative to OCI-Ly7 cells cocultured with BSA beads. P-values (numbers atop of bars) were calculated using one-way ANOVA.

We coated CD40L-tagged TGTs with two force thresholds (12pN and 56pN) on beads, which were incubated with OCI-Ly7 cells within the organoid system (Figure 5A, left). Cells were collected at various time points, permeabilized, stained with antibodies against p-p38 and p-Erk1/2, and analyzed by flow cytometry. It is evident from both fluorescence scattergrams (Figure 5B) and mean fluorescence intensity (Figures 5C and 5D) that at both 2- and 8-h time points, the levels of p-p38 (Figures 5B and 5C) and p-Erk1/2 (Figures 5B and 5D) were elevated when the OCI-Ly7 cells were permitted to exert forces of up to 56pN on the CD40–CD40L bonds, compared to when the cells were restricted from exerting forces beyond 12pN.

### Recording accumulated effect of force through CD40 in vivo by visualizing **α**hCD40-SynNotch activation

To evaluate the in vivo performance of our mechanically activated SynNotch reporter system, we used a luciferase-expressing αhCD40-SynNotch cell line. This was achieved by modifying the previous SynNotch construct’s response element, replacing EGFP with a luciferase enzyme. The luciferase version maintains the same target recognition component, the αhCD40 scFv, and can produce luciferase instead of EGFP upon activation. Due to its higher sensitivity and lower background noise in live mice compared to EGFP, the luciferase assay is particularly suitable for live animal imaging.

Following the establishment of the luciferase version of αhCD40-SynNotch, we encapsulated sender cells, receiver cells, and control cells mixed in various combinations in organoids. Four organoids containing SynNotch expressing Jurkat or plain Jurkat cells and OCI-Ly7 cells were implanted in 4 dorsal subcutaneous (SQ) spaces of each of 6 NOD.Cg-Prkdcscid/J (NSG) mice to ensure adequate statistical power (Figure 6A). The ratio between SynNotch cells and OCI-Ly7 was varied to test sensitivity of SynNotch in vivo while keeping the total cell number constant (10^6^). At various days post-implantation, mice were anesthetized with isoflurane, administered with 3 mg of D-luciferin intraperitoneally, and transferred on a warming bed for bioluminescent imaging for up to 45 min (Figure S6). This was done to assess luciferase reporter activation kinetics at different mixing ratios (Figure 6B). Similar to in vitro results, increasing the proportion of sender relative to receiver cells progressively elevated the signals. Even at a 3:1 ratio, a small number of OCI-Ly7 cells were still capable of activating αhCD40-SynNotch on Jurkat cells, producing significant bioluminescent signals (Figure 6B, blue line). This confirms that CD40 indeed experiences forces sufficient to mechanically activate αhCD40-SynNotch in vivo.

**Figure 6.**
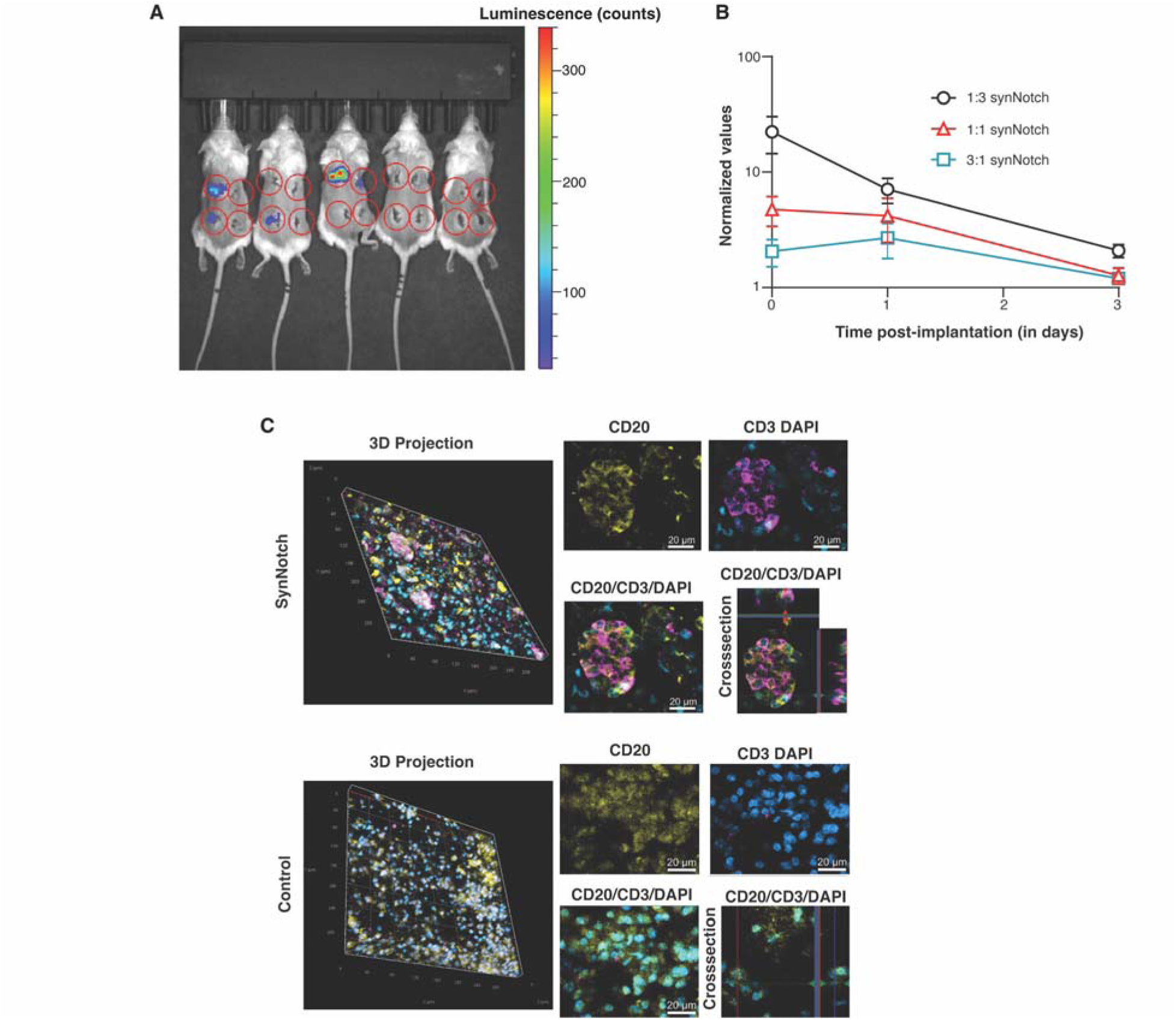
Recording accumulated effect of force through CD40 in vivo by visualizing αhCD40-SynNotch activation. (A) Luminescence images of NSG mice bearing organoids encapsulated with mixture of OCI-Ly7 cells and αhCD40-SynNotch expressing Jurkat cells or plain Jurkat cells. Signal indicates luciferase activity induced by SynNotch activation. Dotted circles represent regions of interest (ROIs) where organoids were implanted. (B) Mean ± SEM of normalized luminescence values (N = 8, n ≥ 10,000) for indicated ratios of SynNotch cells to OCI-Ly7 vs elapsed time post organoid implantation. (C) Representative images of implanted organoids harvested and stained for CD20 (OCI-Ly7), CD4 (Jurkat), and DAPI (both) depicting close contacts between Jurkat cells (purple) expressing (upper group) or not expressing (lower group) αhCD40-SynNotch and OCI-Ly7 cells (yellow) encapsulated in a 1:3 ratio. Images include 3D projections (2 large images on the left column), maximum intensity projections (middle column) and single Z stack/cross-sections (right column).

To further verify that the observed bioluminescence was indeed resulted from luciferase expression upon SynNotch activation, induced by interactions between Jurkat T cells and OCI-Ly7 B cells mediated through CD40–αhCD40-SynNotch bonds, we harvested the implanted organoids, stained for CD20, CD3 and DAPI to identify, respectively, OCI-Ly7, Jurkat, or both cells, and visualized the samples using fluorescence microscopy. We observed that the αhCD40-SynNotch-expressing Jurkat cells formed multiple clusters with OCI-Ly7 cells (Figure 6C, upper), sharply contrasting with the control plain Jurkat cells, which did not form any clusters with OCI-Ly7 cells (Figure 6C, lower).

### In vivo activation of sender cells by receiver cells upon CD40 engagement with **α**hCD40-SynNotch

To determine if B cells are activated by CD40 engagement with αhCD40-SynNotch on sender cells in vivo, organoids were harvested from the dorsal subcutaneous spaces of NSG mice where they were implanted, and the encapsulated cell mixture was released through degradation. B cells were stained with fluorescently conjugated antibodies targeting B cell surface activation markers CD80 and CD86, in addition to phosphorylated intracellular signaling proteins, p-Erk1/2 and p-p38, and analyzed by flow cytometry. The resulting fluorescence scattergrams (Figure S7A) and histograms (Figure S7B) clearly demonstrate that B cells significantly upregulated the expression of CD80, CD86, p-Erk1/2, and p-p38 in comparison to the control groups across the three different receiver-to-sender cell ratios examined. This upregulation is further supported by various quantification methods: mean fluorescence intensity (Figure 7A), their fold change (Figure 7B), the frequency of positive cells (Figure 7C), and the fold changes in these frequencies (Figures 7D and 7E). The data also reveal that the higher the OCI-Ly7 cell number relative to the SynNotch cell number, the higher the levels of CD80/86 expression (Figures 7A and 7B); but the frequency of cells expressing p-Erk1/2 and p-p38 remained relatively constant (Figures 7C-7E).

**Figure 7.**
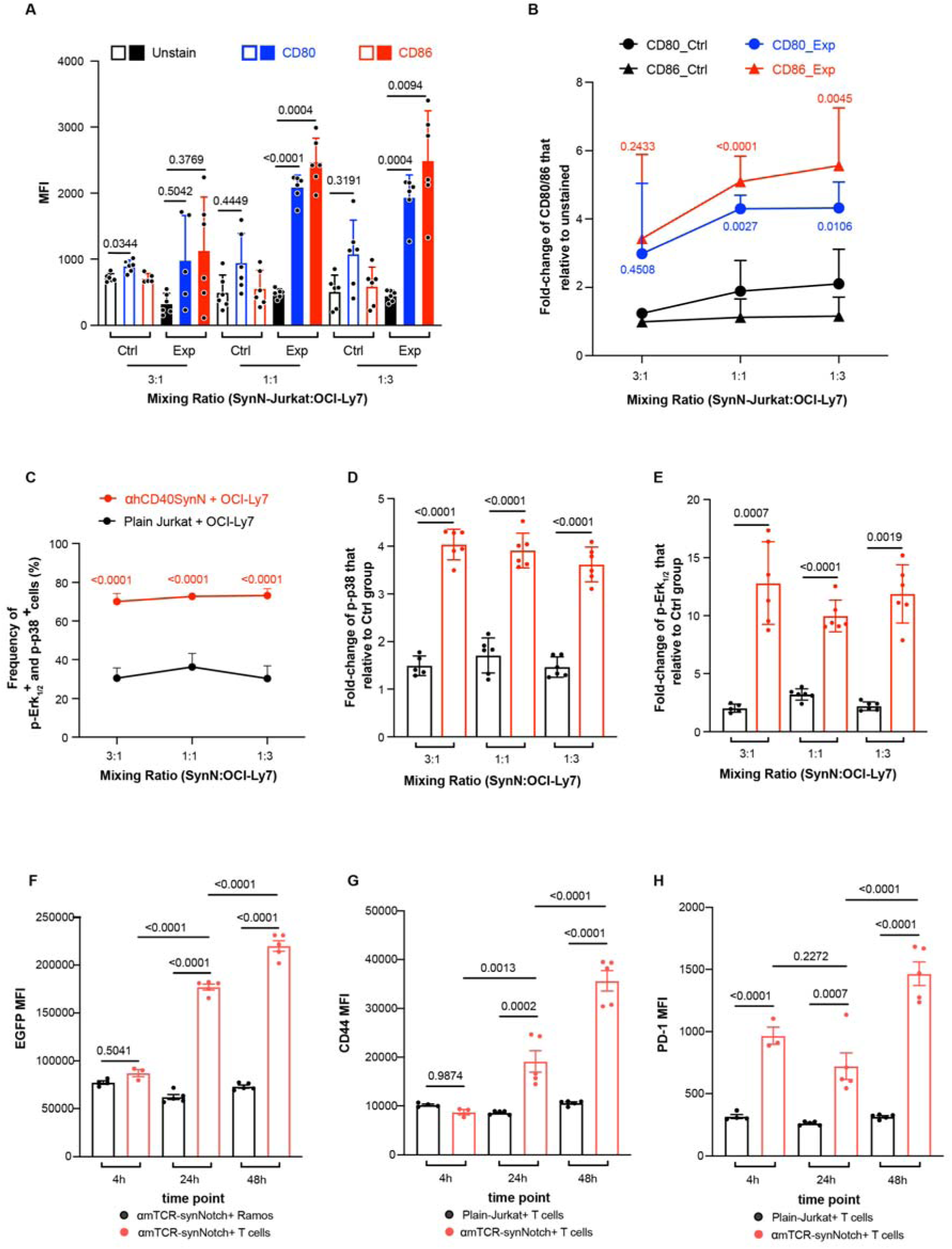
In vivo activation of sender cells by receiver cells upon SynNotch engagement with target receptor. In (A) to (E), OCI-Ly7 cells were harvested and sorted from organoids previously implanted in NSG mice for three days. In (F) to (H), naïve T cells were harvested and sorted from organoids previously implanted in NSG mice for 0, 1 and 2 days. (A) Mean ± SEM with individual data points (# of implants per group N ≥ 5, # of cells n > 10,000) of MFI of fluorescence in sender (OCI-Ly7) cells unstained (black) or stained by antibodies against CD80 (blue) or CD86 (red), which were previously encapsulated with receiver (αhCD40-SynNotch expressing Jurkat) cells (closed) or control (plain Jurkat) cells (open) at indicated ratios. (B) Data in (A) are replotted as mean ± SEM with individual data points of fold-change relative to unstained samples of MFI of CD80 (circle) or CD86 (triangle) staining in OCI-Ly7 cells previously encapsulated in organoids with Jurkat cells expressing (blue or red) or not expressing (black) αhCD40-SynNotch. (C to E) Mean ± SEM (N ≥ 5, n > 10,000) of frequency of p-p38+ and p-Erk1/2+ OCI-Ly7 cells (C) or MFI of p-p38 (D) and p-Erk1/2 (E) staining in OCI-Ly7 cells, which were previously encapsulated in organoids with Jurkat cells expressing (red) or not expressing (black) αhCD40-SynNotch at indicated receiver/sender cell ratios. (F) Mean ± SEM with individual data points (N ≥ 3, n > 10,000) of EGFP MFI in αmTCR-SynNotch expressing Jurkat cells previous encapsulated with naïve T cells expressing mTCR (red) or Ramos B lymphoma cells not expressing mTCR (black) at 1:5 receiver/sender cell ratio at indicated times. (G and H) Mean ± SEM with individual data points (N ≥ 3, n > 10,000) of MFI of CD44 (G) or PD-1 (H) in T cells previously encapsulated in organoids with Jurkat cells expressing (red) or not expressing (black) αmTCR-SynNotch at indicated times. P-values (numbers atop of data) were calculated from one-way ANOVA.

### Recording accumulated effect of force through TCR in vivo by visualizing **α**mTCR-SynNotch activation

We next tested the hypothesis that, in vivo, T cells apply forces on TCRs, and these forces, in turn, activate the T cells. In these experiments, the EGFP version of the αmTCR-SynNotch was used, and 10^5^ Jurkat cells expressing αmTCR-SynNotch mixed with 10^5^ mouse T cells (or Ramos B cells not expressing TCR as negative control) were encapsulated in lymphoid-mimicking hydrogels, which were implanted in NSG mice. At various timepoints post-implantation, hydrogels were harvested from the dorsal subcutaneous spaces of the NSG mice where they were implanted and degraded to release the encapsulated cell mixture. Flow cytometric analysis of Jurkat cells showed significant accumulation of SynNotch activation at 24 and 48 h but not at 4 h. These data indicate that TCR indeed experiences forces transmitted through αmTCR-SynNotch sufficient to mechanically activate reporter expression in this specific in vivo setting (Figure 7F). Concurrently, flow cytometric analysis of T cells co-implanted with αmTCR-SynNotch expressing Jurkat cells found increased CD44 and PD-1 expression over time. Conversely, this upregulation was not observed in T cells co-implanted with controlled Jurkat cells not expressing αmTCR-SynNotch, confirming the dependence of these responses on TCR engagement by αmTCR-SynNotch. These findings were shown by fluorescence scattergrams (Figure S7C), histograms (Figure S7D), and their respective quantifications (Figures 7G and 7H), revealing significant CD44 upregulation at 24 and 48 h post-implantation, as well as significant PD-1 elevation at all three designated time points. These results substantiate our two hypotheses, demonstrating that in vivo, not only do TCRs experience forces, but such forces also act as co-stimuli of T cell activation.

## DISCUSSION

In this work, we demonstrated the construction, characterization, and application of two SynNotch-based mechanically activated transcriptional reporters targeting CD40 and TCR on B and T cells, respectively. We employed two in vitro experiments to test the SynNotch system, which cocultured the receiver and sender cells on a 2D surface and encapsulating them in a 3D hydrogel, respectively, to facilitate physical force transmission through lymphocyte immunoreceptors to mechanically supported bonds with SynNotch molecules, leading to proteolytic activation and downstream reporter expression in receiver cells. After demonstrating the specificity of SynNotch activation, we characterized the sensitivity and reporter activation kinetics of the system. We then used a recently developed PMFA technique to perform a more precise response characterization of SynNotch activation to calibrated externally applied force than a previous report^25^, which defines the force range sufficient to induce downstream reporter expression under the tested conditions. We next used MTP assays to clarify the origin of the force transmission for SynNotch activation, verifying that SynNotch-expressing receiver cells could not generate detectable endogenous force required for reporter activation and confirming that receptor-expressing sender cells generated forces on the receptor–SynNotch bonds. This allows us to attribute the reporter signal to force generated by the sender cells and exerted on the target receptor upon engagement with the SynNotch, thereby placing the application of our system on a more rigorous foundation. We have recently reported that primary B cells and a B lymphoma cell line (Farage) exert endogenous forces to bear on CD40–CD40L bonds, using three MTPs with threshold forces of 4.7, 12, and 19pN to pinpoint the force level to be ∼15 pN^16^. Since our calibration curve reveals that SynNotch reporter is activatable by forces as low as 2.4pN, we used the 4.7-pN MTP in the present study to achieve the maximum force sensitivity, allowing us to conclude that receiver cells are unable to exert >4.7pN forces on αhCD40-SynNotch at least during the first 5-h of receptor engagement.

Through coculturing receiver and sender cells in both 2D and 3D conditions, our in vitro experiments confirmed activation of both pairs of the SynNotch reporters and the target receptors, thereby demonstrating that forces are transmitted through CD40 and TCR bonds bridging cell-cell junctions, not solely between cells and inert surfaces, thus addressing a controversy within the field. However, because SynNotch activation is a process involving irreversible cleavage followed by reporter transcription and translation, the eventual output should be interpreted as an accumulated readout of SynNotch activation rather than an instantaneous measurement of force magnitude. The temporal delay from force transmission to reporter detection may result from the fact that mechanical inputs at receptor–ligand bonds likely occur on much shorter timescales than the hours required for the biological process that involves SynNotch TMD cleavage, nuclear translocation of the released intracellular domain, transcription, translation, and EGFP maturation. During the 3-5 h window before detectable reporter signal emerges, other biological processes may also occur, which may or may not depend on force, including receptor trafficking, cytoskeletal remodeling, changes in gene expression, and reporter maturation. In addition, reporter signal reflects the net balance between reporter production, maturation, dilution, and degradation. Therefore, the onset, plateau, or decline of EGFP or luciferase signal should not be interpreted as direct evidence for the onset, persistence, cessation, or magnitude of mechanical force. Instead, the time courses in our experiments represent time-integrated reporter kinetics following mechanically activated SynNotch cleavage and signaling.

To demonstrate potential in vivo applications, we implanted organoids encapsulating receiver/sender cell mixtures into NSG mice. Our dual observations of SynNotch activation in receiver cells and target receptor signaling in sender cells provide in vivo evidence supporting two key conclusions: (1) B and T cells indeed generate endogenous forces to exert on CD40 and TCR, respectively; and (2) such forces are biologically relevant as they reinforce the activation of B and T cells. The second conclusion addresses another controversy in the field. A caveat is that our in vivo experiments lasted 1-3 days during which time the receiver cells might also exert migration-generated force on receptor–SynNotch bonds to activate reporter expression, as observed in the case of αmTCR-SynNotch. This prevents us from attributing the reporter signals solely to endogenous forces generated by sender cells. The NSG mouse also suffers key limitations because it is immunodeficient, which lacks functional T, B, and NK cells, hence contributing no adaptive immune component. The subcutaneous space where our encapsulated sender and receiver cells were implanted also provides no lymphoid architecture. Future studies using immunocompetent settings will be important to determine how this reporter system performs in more physiological immune environments. These should also utilize more pertinent disease animal models to examine the mechanical environment of various target receptors on diverse host cells, which are not encapsulated with the receiver cells but are recruited through SynNotch interactions with the target receptors.

In contrast to MTP, TFM, and mPADs, which utilize inert materials to construct a “strain gauge” for force measurement, the SynNotch system leverages the intrinsic properties of a biological molecule to develop a delayed force reporter. Our data confirm that engagement of the target receptor does not cause the receiver cell to exert force on SynNotch, thereby supporting the interpretation that the reporter readout primarily reflects mechanical input from sender cells rather than forces originating from the receiver cells, a characteristic shared by the existing methodologies. Like the existing techniques, the SynNotch system is repurposed for detecting forces from target receptors on living cells, which would not exert force if the ligands or antibodies engaged on the target receptors are soluble due to the lack of mechanical support to counter-balance the reaction force^51^. Regardless of whether the forces through immunoreceptors were generated by sender cells or due to receiver cell migration, several requirements for detecting force signals can be envisioned. Initially, sensor must bind target and such binding must be maintained for a prolonged duration, which reflects the significance of binding affinity and bond lifetime. In our study, these two criteria have been met by employing an antibody scFv directed against the target receptor of interest whose affinity and force-dependent bond lifetime were measured. The observed SynNotch reporter activation in our system indicates at least a portion of receptor–SynNotch bonds are mechanically sustained, or repeatedly rupture and rebind, for sufficient durations to permit trans-activation. Subsequently, receptor engagement must initiate signaling, as the sender cell needs to activate its force generation and transmission mechanisms. Furthermore, the process from signal initiation to the application of force should incorporate feedback mechanisms and mechanical reinforcement, enabling the sender cell to differentiate between binding of immobilized and soluble ligands to its receptor. Indeed, by replacing the sender cells with beads functionalized with TGT-tagged CD40L, we demonstrated not only receptor-mediated signaling in the sender cell but also the signal amplifying effect of endogenous force on CD40. This data is consistent with our recent report that permitting up to 56pN of endogenous forces on CD40–CD40L bonds induces higher signaling of primary B cells and a B lymphoma cell line (Farage) than preventing such force from exceeding 12 pN^16^.

It should be noted that the second and third requirements—namely, sender cell signaling induced by receptor binding and signal amplification through mechanical reinforcement—are of particular significance to the field of mechanobiology, as they are fundamental to receptor-mediated mechanotransduction. In the TCR case, they underpin key premises of the mechanosensor model and represent longstanding questions related to TCR signal initiation and antigen discrimination, which are essential to T cell adaptive immunity. By using the mechanically activated SynNotch, our work addresses these questions and supports these premises, contributing substantial evidence for the mechanosensor model.

## METHODS

### Cultured cells

Jurkat T cells (clone E6.1) purchased from American Type Culture Collection (ATCC) were used as receiver cells to express the four SynNotch molecules. P40.2B4 cells and Ramos cells were generous gifts of Dr. Michelle Krogsgaard (New York University, New York, NY, USA), Dr. Jürgen Wienands (University Medical Center Göttingen, Göttingen, Germany), respectively, were used as sender cells or control cells. OCI-LY7 were originally provided by the University Health Network, Toronto, and transferred by scientist Dr. Mark Minden. HEK293T/17 cells, also from ATCC, were used to produce SynNotch-encoding Lentiviruses. All cells were cultured at 37 °C with 5% CO2. HEK293T/17 cells were cultured in Dulbecco’s modified Eagle’s medium (DMEM) high glucose (4.5 g/L), with supplements of L-glutamine (6 mM), MEM Non-Essential Amino Acids (NEAA, 0.1 mM), Sodium Pyruvate (1mM), Fetal Bovine Serum (FBS) (10%). Jurkat cells and 2B4 cells were cultured in R10 medium, consisting of RPMI 1640 supplemented with FBS (10%), Penicillin-Streptomycin (1%), HEPES (10 mM) and Sodium Pyruvate (1 mM). Ramos cells were cultured in RPMI 1640, with supplements of FBS (10%), penicillin-streptomycin (1%), HEPES(10mM), L-glutamine (1 mM), sodium pyruvate (1 mM), and β-mercaptoethanol (50 μM). OCI-Ly7 cells were cultured in IMDM with supplements of FBS (10%), Penicillin-Streptomycin (1%), L-glutamine (1 mM), MEM Non-Essential Amino Acids (NEAA, 0.1 mM).

### Primary cells and mice

Peripheral blood mononuclear cell (PBMC)-derived B cells were purified from freshly isolated blood of healthy donors according to a protocol approved by the Institutional Review Board (IRB) of the Georgia Institute of Technology (protocol No. H22021). Briefly, after diluting whole blood 1:1 with PBS, PBMCs were separated by density gradient centrifugation using Histopaque-1077 (Millipore Sigma) and subjected to low-speed centrifugation washes (120 × g). CD19+ B cells were subsequently purified from the PBMC fraction using the EasySep Human CD19 Positive Selection Kit II (STEMCELL Technologies). After purification, the B cells were kept in RPMI supplemented with 10% FBS, 1% penicillin-streptomycin, 10 mM HEPES, and 1× nonessential amino acids, and used immediately for each experiment. Naïve CD8^+^ mouse primary T cells were used as sender cells for the αmTCR-SynNotch expressing receiver cells. They were isolated from the spleen of either P14 or OT-I transgenic mice with untouched CD8^+^ T cell isolation kit (Stemcell Technologies).

P14, OT-I, and NOD.Cg-Prkdcscid/J (NSG) mice were housed either at the Emory University Department of Animal Resources facility following protocol approved by the Institutional Animal Care and Use Committee of Emory University or at the Georgia Tech Animal Facility following protocol ZHU-A100168-07/26/2025 and Kwong-A A100190 approved by the Institutional Animal Care and Use Committee (IACUC) of Georgia Tech.

### Generation of stable expressions of SynNotch and response element

The response element plasmid containing a PGK promoter, which drives the constitutive expression of a blue fluorescent protein (BFP) or mCherry, and the original antiCD19-SynNotch construct containing a Myc tag are gifts from Wendell Lim (University of California, San Francisco, CA, USA). The luciferase version of the reporter was provided by the Kwong lab and was derived from the Addgene pHR_Gal4UAS_pGK_mCherry response-element plasmid deposited by the Lim lab. The construct retained the Gal4UAS response-element architecture, with an additional luciferase-encoding sequence inserted into the plasmid. To construct the αhCD40SynNotch and αmTCRSynNotch plasmids, the ligand binding antiCD19-scFv in original SynNotch construct containing a Myc tag was replaced by the scFv of an anti-human CD40 (derived from the C224S/kC214S IgG2 variant^52^, PDB: 6TKE) or the anti-mouse TCR antibody clone H57 (a gift from Johannes Huppa of Charité – Universitätsmedizin Berlin, Germany). The SynNotch and Gal4VP64 trans-activation plasmids contained an ampicillin resistance gene (Amp^R), which was used for bacterial selection in competent cells on LB agar plates containing 100 µg/ml ampicillin.

Lentiviral particles were produced in HEK293T/17 cells by co-transfecting with package and envelop plasmids using Lipofectamine 3000. Viral supernatants were collected at 72 h post-transfection, cleared by centrifugation and concentrated using Lenti-X Concentrator according to the manufacturer’s instructions.

For transduction, Jurkat cells were resuspended in medium containing concentrated lentivirus with the response element and SynNotch with αhCD40 or αmTCR svFv and 10ug/ml polybrene. Cells were then spin-transduced by centrifugation at 1200 × g for 1h at 30 °C, with both acceleration and deceleration set to 0. transduced with plasmids containing the response element and SynNotch or with αhCD40 or αmTCR svFv sequentially. To generate stable cell lines expressing desired expression levels of transduced genes, cells went through multiple rounds of FACS sorting for uniform levels of BFP/mcherry (marker for the response element) and anti-Myc antibody conjugated with PE/AF647 (marker of SynNotch).

### Proteins, antibodies, and chemicals

C-terminally biotinylated soluble human CD40 protein was produced in-house^16^. C-terminally biotinylated soluble mouse OT-I TCR protein was a kind gift of David Margulies^53^ (National Institute of Allergy and Infectious Diseases, NIH, Bethesda, MD, USA). The antibodies, chemicals and other reagents are summarized in Supplementary Table 1.

### Flow cytometry

Flow cytometry was used to quantify SynNotch expression and activation in receiver cells as well as target receptor expression and triggering (or the lack thereof) on sender cells (or control cells). Samples (cells or beads) were collected and washed at 4 °C with FACS buffer (PBS without Ca^2+^ or Mg^2+^, 5mM EDTA, 2% FBS) two times. Samples were then stained for 30 min at 4 °C in 100 μl of FACS buffer containing 10 μg/ml (or as suggested by the manufacturer’s instruction) of antibodies against the Myc tag (for SynNotch expression), CD40 and TCR (for target receptor expression), p-p38 and p-Erk1/2 as well as CD80 and CD86 (for CD40 signaling-induced B cell activation), and CD44 and PD-1 (for TCR signaling-induced T cell activation), washed twice with 1 ml of FACS buffer, and fixed with 300 μl of 4% PFA for 15 min at 4 °C. Samples were washed once with 1 ml of FACS buffer and resuspended in 300 μl of FACS buffer for analysis under BD Fortessa (BD Biosciences) or Cytek Aurora (Cytek Biosciences). For samples loaded in the 96-well plate, Cytoflex (Beckman Coulter) was used to allow high-throughput readout. Flow cytometry data were analyzed using FACS DIVA (BD Biosciences). For determination of site densities of CD40 and αhCD40SynNotch for 2D affinity measurement, standard beads were included in the samples to produce respective calibration curves^16^. FlowJo software (Treestar) was used for analysis of all flow cytometry data in this study.

### Micropipette adhesion frequency assay

To measure effective 2D affinity between CD40 and αhCD40-SynNotch, we used the micropipette adhesion frequency assay, which has been thoroughly described^16,42^. In brief, red blood cells (RBCs) were biotinylated using EZ-Link Sulfo-NHS-LC-Biotin, incubated with streptavidin, and then with C-terminally biotinylated soluble CD40 protein. Functionalized RBCs and αhCD40-SynNotch expressing Jurkat cells were loaded into a chamber on opposite sides to avoid mixing. The chamber was then mounted onto an inverted microscope and glass micropipettes were inserted into the chamber. An RBC and Jurkat cell pairs were aspirated onto opposite pipette tips with minimal pressure. Using a computer-controlled piezo translator attached to one pipette, cells were brought into repeated contacts for a 2-s duration and constant area for each contact. Binding was visualized microscopically and enumerated for a total of 50 touches for each individual cell pair to determine an adhesion frequency *P*_a_. The average number of bonds was calculated by <*n*> = - ln(1 – *P*_a_). The effective 2D affinity was calculated using the site densities of the target receptor CD40 (*m*_r_) and αhCD40-SynNotch (*m*_l_) on the RBC and Jurkat cells (measured by flow cytometry using standard beads) by *A*_c_*K*_a_ = <*n*>/(*m*_r_*m*_l_). Using the values of *A*_c_*K*_a_ and *m*_l_, we can then calculate the <*n*> values between αhCD40-SynNotch expressing Jurkat cells and paramagnetic beads coated with a range of CD40 densities in the PMFA experiment to generate the force response curve for SynNotch activation.

### Force-dependent bond lifetime measurement by biomembrane force probe

To generate a force response curve for SynNotch activation also requires information regarding how force modulates dissociation of SynNotch scFv from the target receptor, which was measured using a biomembrane force probe. Our custom-design and home-made biomembrane force probe has been described^16,54,55^. In brief, the biomembrane force probe uses a bead attached to the pressurized RBC as a force transducer to enable optical tracking of its position with high spatial (a few nanometers) and temporal (< 1ms) resolution. Force as small as a single digit piconewton is determined from the deflection of the RBC membrane. Beads and cells were prepared similarly as the micropipette adhesion frequency assay. To ensure that most adhesion events were mediated by a single molecular bond, adhesion events were controlled to be infrequent (≤ 20%). In the retraction phase of the mechanical cycle, if a bond survived the ramping and reached a preset tension level, the force was clamped until spontaneous bond dissociation and a pair of values of the clamped force *f* and bond lifetime *t*_b_ were measured. Bond lifetimes were measured at forces ranging from 2 – 40pN, pooled, and binned into >5 force bins (>35 measurements per bin) to reduce system errors, and presented as mean bond lifetime 〈*t*_b_〉 ± SEM.

### Parallel magnetic force activation (PMFA) assay

The design of PMFA assay has been described^16^. First, we applied a droplet of epoxy using a pipette tip to the center of each well (∼5-mm area) of a non-tissue culture treated 24-well plate. At the same time, 5 mm-diameter glass coverslips were cleaned by sequential washes in 70% EtOH and di-H_2_O. Each coverslip was dried by a kimwipe tissue and gently placed on top of the epoxy droplet. After attaching coverslips, the wells were washed with 1 ml PBS and incubated in 1 ml PBS for 30 min at room temperature. After PBS was thoroughly aspirated and air dry completely, a 18μl droplet of 0.01% PLL solution was carefully added to each coverslip to incubate at 4 °C overnight. On the following day, wells were washed with PBS in the same manner, followed by addition of 18μl of cells (1-2 x 10^5^) in culture medium on top of each coverslip and incubation for 10 min at 37 °C to immobilize cells. Then 2μl of a well-mixed CD40-coated paramagnetic beads in culture medium were carefully added on top of the R10 droplet above the immobilized cells for 10 min incubation at 37 °C to allow beads to settle on the cells by gravity. An additional 80μl of R10 medium was added to each well around the coverslip for 10 min incubation at 37 °C. The lid with magnets (or without for control samples) was gently placed on top of each plate. After designated time of incubation, cells were harvested and analyzed EGFP expression by flow cytometry.

### Molecular tension probe (MTP) assay

To prepare the MTP surface, 25 mm glass coverslips were sonicated in 50% ethanol for 15 min and rinsed 3X with di-H_2_O. Following rinsing, the coverslips were immersed in 40 ml Piranha solution (mixture of 25 ml sulfuric acid and 15 ml H_2_O_2_) for 30 min. Coverslips were washed 6X with di-H_2_O and 3X with 100% ethanol. The surfaces were then salinized with 3% APTES in 200 proof ethanol for 1 h, then washed 3X in ethanol and dried at 80°C for 30 min. After drying, 200μl of 10 mg/ml LA-PEG (Biochempeg), 50 mg/ml mPEG (Biochempeg) in 0.1 M NaHCO_3_ was added to each surface and incubated for 1 h at room temperature. Surfaces were then washed 3X with di-H_2_O and 200μl of 1 mg/ml Sulfo-NHS-Acetate was added to two coverslips placed together as a sandwich and incubated for 30 min at room temperature. Coverslips were again rinsed 3X with di-H_2_O and 500μl gold nanoparticle (Au-NP, 8.6 ± 0.6 nm diameter) solution was added to each sandwich and incubated for 30 min at room temperature. During Au-NP incubation, 300nM hairpin strand, 330nM quencher strand (BHQ2), and 330nM Cy3b strand of ssDNA in 1 M NaCl were annealed in a thermocycler by heating to 95°C for 5 min and gradually cooling (−5°C/min) to 25°C. After rinsing Au-NP coated surfaces with 3X di-H_2_O and 2X 1 M NaCl washes, 100μl of annealed DNA were added to each sandwich and incubated overnight at 4°C. The following day, the coverslips were rinsed with 3X PBS after which 40μg/ml streptavidin in PBS was added to each sandwich and incubated for 1 h at room temperature. Following 3X rinse with PBS, 40μg/ml of C-terminally biotinylated soluble human CD40, mouse OT-I TCR, anti-CD3 antibody (positive control), or BSA (negative control) in PBS + 2% BSA was added to each sandwich and incubated for 1 h at room temperature.

To perform MTP experiment, cells were loaded in the imaging chamber (formed by the MTP-functionalized coverslip as the chamber floor) and incubated for ∼30 min (except one experiment in which a prolonged incubation of ∼5 h was used). Cells were imaged using Nikon W1 spinning disk confocal microscope equipped with a Plan-Apochromat 60x/1.40 oil objective. The RICM module was used to detect the cell bottom and measure spreading area, the TIRF mode was used to measure the Cy3b fluorescence, and the epi-fluorescence mode was used to quantify EGFP fluorescence.

### Time-lapse SynNotch activation using a 2D coculture system

Receiver cells (Jurkat) expressing SynNotch with scFv of either anti-human CD40 or anti-mouse TCR and sender cells (OCI-Ly7 or naïve CD8^+^ T cells experimental group, 2B4 or OCI-Ly7 as negative control, depending on the scFv used) were counted and mixed in 1:3 or 1:5 SynNotch:Sender ratio. Cells were resuspended in 150μL culture medium in U-shape 96-well plate and spin-down using centrifuge (100 g, 1 min) to provide cell-cell contact. Different starting points were chosen to obtain SynNotch time-lapsed activation profiles. At end point, cells were fixed and their EGFP expressing level was checked using Cytoflex platform (Beckman Coulter).

### Time-lapse SynNotch activation in a 3D organoid system

The receiver and sender cells were encapsulated in hydrogel-based organoids following the previously described protocol, similar to the 2D coculture experiment^46^. Briefly, hydrogel-based organoids were composed of 4 arm Maleimide functionalized polyethylene glycol (PEG-4-MAL) functionalized with integrin specific peptides and crosslinked with matrix metalloproteinase (MMP) degradable, VPM (GCRDVPM↓SMRGGDRCG peptide), and nondegradable, Dithiothreitol (DTT), peptides. VPM and DTT were incorporated at 1:1 ratio^46^. 7.5 wt% PEG-4MAL organoids were functionalized with RGD oligopeptide (GRGDSPC), fibronectin mimetic that binds to αVβ3 integrin^46^.

OCI-LY7 cells were originally provided by the University Health Network, Toronto, and transferring scientist Dr. Mark Minden. OCI-Ly7 cells and αhCD40-SynNotch-expressing Jurkat cells were encapsulated at a 1:3 ratio (50,000 Jurkat cells: 150,000 OCI-Ly7 cells) and cultured at various timepoints (0, 1, 3, 6, 12, 24, 48 and 96 hours). Naïve OT-1 CD8^+^ T cells and αmTCR-SynNotch-expressing Jurkat cells were encapsulated at a 1:1 ratio (100,000 Jurkat cells: 100,000 T cells) and cultured for 48 hours. At the end point, organoids were enzymatically degraded using collagenase (Fisher Scientific, NC9482366) for 1 h. The resulting cell suspensions were washed twice with PBS buffer.

For analysis of the αhCD40-SynNotch system, half of the cell suspensions were stained at 4 °C for 30 min in 100μl of FACS buffer containing 1μl of anti-CD20-BUV805, anti-CD80-PE, and anti-CD86-APC antibodies (Fisher Scientific). Samples were subsequently washed twice with 200μl of FACS buffer, fixed in 100μl of 4% paraformaldehyde (PFA) for 20 min at room temperature, washed once more with 200μl of FACS buffer, and resuspended in 100μl of FACS buffer for flow cytometry analysis using a CytoFLEX system (Beckman Coulter). The remaining half of the samples were fixed in 100μl of 4% PFA for 20 min at room temperature, followed by centrifugation at 600 g for 5 min at 4 °C (acceleration: 9, deceleration: 9). Cells were then resuspended in 200μl of methanol at 4 °C for 30 min, washed with 200μl of FACS buffer, and stained with anti-CD20-BUV805, anti-Human/Mouse (Hu/Mo) Phospho-p38-APC, and anti-Hu/Mo Phospho-ERK1/2-PE antibodies (Fisher Scientific) at 4 °C for 30 min. Samples were then washed, resuspended in FACS buffer, and analyzed by flow cytometry.

For analysis of the αmTCR-SynNotch system, all cell suspensions were stained at 4 °C for 30 min in 100 μl of FACS buffer containing 1μl of anti-CD8-PE (biolegend), anti-CD44-APC (Cytek Biosciences), and anti-PD1-Alexa Fluor 750 antibodies (Bio-Techne). Samples were subsequently washed twice with 200μl of FACS buffer, and resuspended in 100μl of FACS buffer for flow cytometry analysis using a Cytek Aurora (Cytek Biosciences).

### DAPT inhibition experiment

DAPT was dissolved in cell-culture grade DMSO to a stock concentration of 10 mM and diluted in cell culture media to 10 μM. DAPT-containing media or DMSO control media were added to αhCD40-SynNotch-expressing Jurkat and incubated for 4 h at 37°C. 3×10^4^ SynNotch cells were mixed with 7×10^4^ primary human B cells and encapsulated in each organoid. DAPT and DMSO were maintained in the culture medium throughout the experiment. Cell mixtures in organoids were cultured for 12 and 24 h followed by enzymatic degradation to recover cells and analysis by flow cytometry for EGFP expression. αhCD40-SynNotch-expressing Jurkat cells encapsulated with 2B4 cells not expressing CD40 and plain Jurkat cells not expressing αhCD40-SynNotch encapsulated with primary B cells were used as negative controls.

### Tension gauge tether (TGT) assay

Tension gauge tether (TGT) probes were annealed using the same conditions described above for the molecular tension probe assay. Briefly, each DNA strand was mixed at 220nM in 1M NaCl in a 1 mL reaction volume and annealed in a thermocycler). Following annealing, azide-coated PMMA beads (12μm, PolyAn) were prepared at a 1:1 cell-to-bead ratio and washed twice in PBS + 2% BSA by centrifugation at 200g for 3 minutes. The beads were then resuspended in TGT probe solution (∼1 mL) and incubated overnight at room temperature under continuous rotation to allow covalent attachment of the TGT probes to the bead surface through strain-promoted azide-alkyne cycloaddition (SPAAC) click chemistry between the azide groups on the beads and the dibenzocyclooctyne (DBCO)-modified TGT oligonucleotide. In the following day, TGT-coated beads were washed twice with PBS + 2% BSA, resuspended in 500μl of 40μg/ml streptavidin (PBS + 2% BSA), and incubated for 1 h at room temperature under rotation. Beads were then washed twice, resuspended in 1μg/ml biotinylated CD40L (PBS + 2% BSA), and incubated for 1 h at room temperature. Excess CD40L was removed by washing twice in PBS + 2% BSA before use.

To use TGT to study the effect of limiting CD40 force on B cell activation, OCI-Ly7 cells and TGT-coated beads were encapsulated in organoids at a 1:1 ratio (100,000 TGT beads: 100,000 OCI-Ly7 cells) and cultured at various time points (0, 2, 4, 6, 8, and 10 hours). Organoids were then enzymatically degraded with collagenase for 1 h, followed by two PBS washes. Cells were fixed in 100μl of 4% PFA for 20 min at room temperature, followed by centrifugation at 600 g for 5 min at 4°C (acceleration: 9, deceleration: 9). The samples were then resuspended in 200μl of methanol at 4 °C for 30 min, washed with 200μl of FACS buffer, and stained with anti-Hu/Mo Phospho-p38-APC and anti-Hu/Mo Phospho-ERK1/2-PE antibodies (Fisher Scientific) at 4 °C for 30 min. Following washing and resuspension in FACS buffer, samples were analyzed by flow cytometry using a CytoFLEX system.

### Surgical hydrogel implantation into the subcutaneous space

All animal procedures were conducted as approved by the Institutional Animal Care and Use Committee (IACUC; protocol number A100368) and followed the guidelines mentioned in the Guide for the Care and Use of Laboratory Animals. Hydrogels were prepared as discussed earlier and implanted in the dorsal subcutaneous (SQ) space of mice using a minor surgery^56^. Briefly, male NOD.Cg-Prkdcscid/J mice were anesthetized under isoflurane (3% v/v for induction, followed by 1.5-2% v/v for maintenance) and transferred to a warming bed to undergo the surgical procedure. For this duration, the mice were in a prone position and anesthesia was administered using a nose cone. Fur from the dorsal region was shaved and the skin was sanitized with 70% ethanol. A small incision was made in the skin and the underlying connective tissue cleared using blunt forceps, thus resulting in an SQ pocket. Pre-cast hydrogels (maintained in media from the time of preparation to the time of implantation; N=5-8 hydrogels per group) containing cells in the desired ratio were implanted into this space, followed by wound closure using 4-0 sutures. This procedure was performed in a single-blind technique (surgeon was blinded) for up to 4 implants per mouse. The position of implants in the dorsal SQ space was randomized to remove any experimenter bias. To manage post-surgical pain, animals were administered buprenorphine-SR (1 mg/kg) and monitored closely for signs of distress.

### In vivo imaging by IVIS spectrum

We evaluated the successful activation of anti-hCD40SynNotch signaling in implanted hydrogels by quantifying their bioluminescent activity. Briefly, mice were anesthetized at predetermined time-points as mentioned earlier and were injected with D-luciferin (3 mg) intraperitoneally. The animals were then transferred to imaging chamber of the PerkinElmer IVIS Spectrum CT where bioluminescent images were acquired up to 45 min after injection. Images were subsequently analyzed for bioluminescent signal intensity using LivingImage® 4.7.4 software.

### Analysis of in vivo activation by flow cytometry

To evaluate the in vivo activation of αmTCR-SynNotch and TCR-mediated T cell signaling in hydrogels implanted in NSG mice, mice received SQ implants as detailed above (up to four implants per mouse). At respective durations post-implantation, mice were euthanized and their implants collected. The implants were enzymatically degraded (collagenase I, 200 U/mL, at 37°C for up to 30 mins) to release the cells of interest. the resulting cell suspensions were washed twice with FACS buffer and stained at 4°C for 30 min in 100μl of FACS buffer containing 1 ul of anti-CD8-PE (Fisher Scientific), anti-CD44-APC (Cytek Biosciences), and anti-PD1-Alexa Fluor 750 antibodies (Bio-Techne). Samples were subsequently washed twice with 200μl of FACS buffer, and resuspended in 100μl of FACS buffer for flow cytometry analysis using a Cytek Aurora (Cytek Biosciences).

### Statistics

Results are presented as mean ± SEM, or as mean values only when appropriate. Statistical analyses were performed using GraphPad Prism. The choice of statistical comparisons was determined based on data type, distribution, and study design. Methods used include the Student’s t-test and one-way ANOVA. Custom LabView and Julia scripts were developed to analyze BFP and MTP datasets. Data obtained from flow cytometry, TGT, and PMFA assays were processed and analyzed using FlowJo and MATLAB.

## Acknowledgements

This work was supported by NIH grants U01CA250040 (C.Z.), U01CA280984 (A.S. and C.Z.), R01CA238745 (A.S.), and HD091793 (G.A.K.). H.-K.C. was partly supported by a National Research Foundation grant of South Korea (2021R1A6A3A03038382) and D.B was supported by ImmunoEngineering training Grant 2T32EB021962-06A1. A.S. acknowledges support from the Carl Ring Family Professorship. We thank Wendell Lim for the αCD19-SynNotch plasmid. We also thank Michelle Krogsgaard for providing P40.2B4 cells, Jürgen Wienands for providing Ramos cells, and Ari Melnick for providing OCI-Ly7 cells. We thank David Margulies for providing soluble TCR protein and Stefano Travaglino for making soluble CD40 protein. We thank Peiwen Cong, Larissa Doudy, and Matthew Wang for technical assistance.

## Author contributions statement

J.L., M.L, K.L., A.S., and C.Z designed experiments; M.L., J.L., A.D., D.B. and A.A. performed experiments; M.L., J.L., A.D. and A.A. analyzed the data. G.A.K. contributed key reagents. M.L, J.L. and C.Z. wrote the initial draft and all other authors contributed to revising and finalizing the manuscript. A.S and C.Z secured funding.

## Data and materials availability

All data are available in the main text or the supplementary materials.

## Supplementary Materials

Figs. S1 to S7 Table S1

