## Supplementary Information for "SynNotch receptors for visualizing immunoreceptor force transmission and downstream signaling in vivo"

**The PDF file includes:**

Figs. S1 to S7

Tables S1

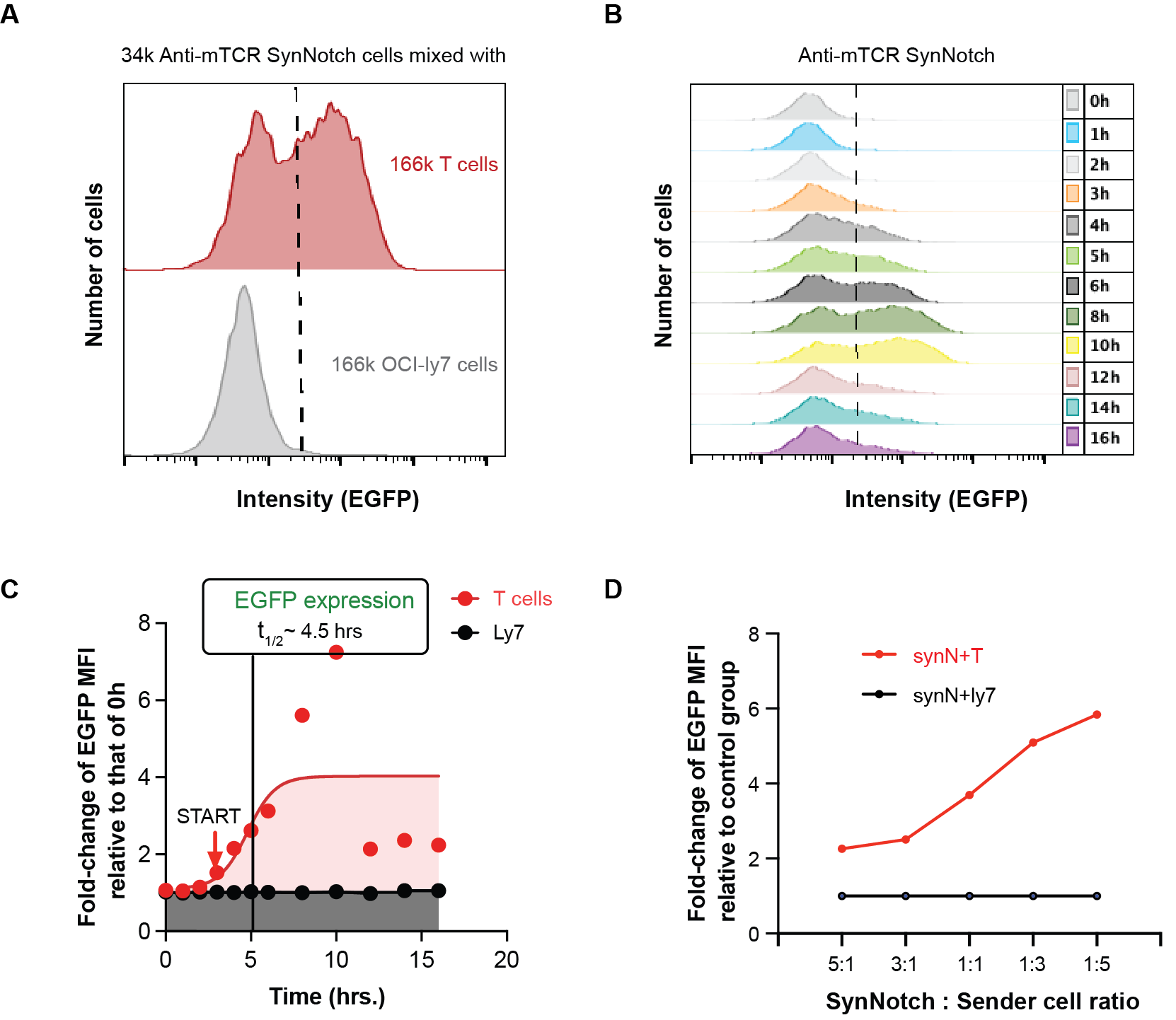

Fig. S1. Reporter activation kinetics of αmTCR-SynNotch in a 2D coculture system. (A) Histograms of EGFP fluorescence in receiver cells (Jurkat) expressing αmTCR-SynNotch cocultured with 5X excessive sender cells (naïve CD8^+^ mouse T cells) expressing TCR or control cells (OCI-ly7) not expressing the target receptor, confirming the specificity of αmTCR-SynNotch activation. (B) Representative fluorescence histogram stack of EGFP expression in αmTCR-SynNotch expressing Jurkat cells at various times. The 96-well plate layout and experimental procedure were the same as Fig. 1C except that the sender and control cells were changed to T and B cells, respectively, and the SynNotch scFv on the receiver cells (3.4×10^4^) was changed from anti-hCD40 to anti-mTCR. (C) Mean ± SEM (N = 2, n > 10,000) of fold-change of EGFP MFI (point) and sigmoidal model fit (curve) of αmTCR-SynNotch expressing Jurkat cells cocultured with mouse T cells cells (red) or OCI-Ly7 not expressing TCR (black) at various times relative to that of 0 h. Red arrow with START indicates the starting point of EGFP upregulation relative to that of 0 h (N = 2, n > 10,000) at various of time points. (D) Data points (N = 1, n > 33,000) of fold-change of EGFP MFI relative to that of SynNotch only condition at different ratios for 8 hours.

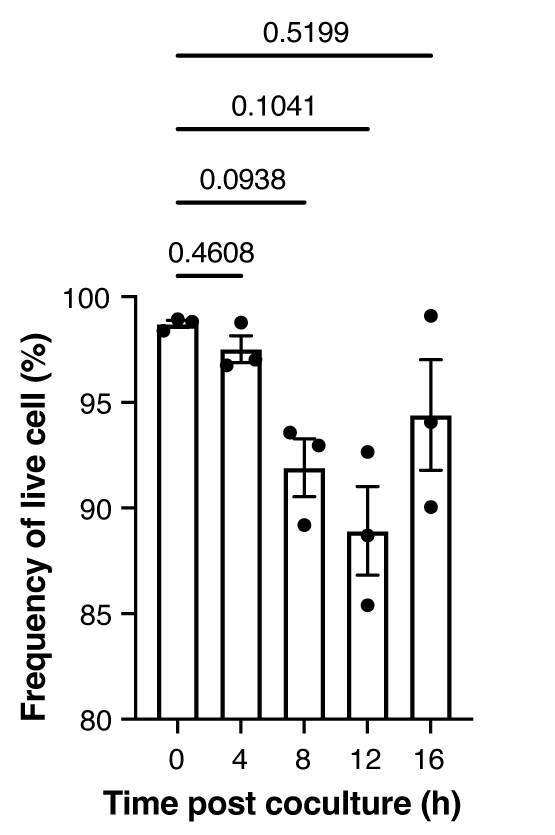

Fig. S2. Cell viability and αhCD40-SynNotch expression level. Mean ± SEM (plus individual data points of three repeated experiments) of frequency of live cells. P-values were determined using one-way ANOVA.

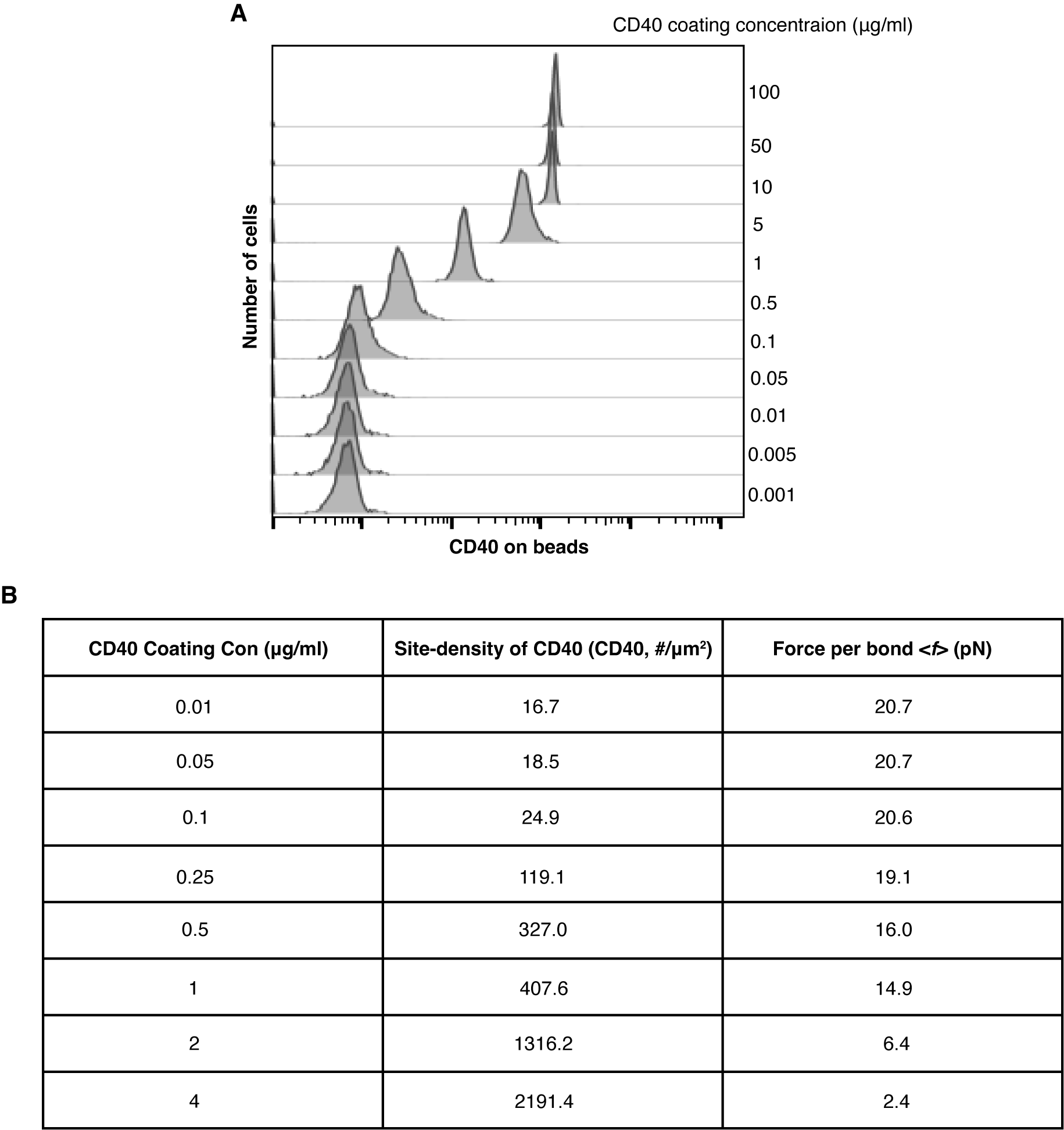

**Fig. S3.** **Relating coating concentration of soluble CD40 to its site density on paramagnetic bead and average force per bond**. **(A)** Representative histogram stack of fluorescence staining by an anti-CD40 antibody on 2.8-μm paramagnetic beads at various coating concentration of soluble CD40 protein. **(B)** Table showing CD40 coating concentration (left column) with corresponding site-density of CD40 coated on paramagnetic beads (middle column) and average force per bond (middle column), calculated by

$$\left\langle f \right\rangle=\frac{F}{\exp\left( \left\langle n \right\rangle\right)-1}\sum_{n=1}^{\infty} \frac{\left\langle n \right\rangle^{n}}{nn!}$$

where *F* is the magnetic force per bead determined in the PMFA design(*16*) and the average number of bonds per contact is

$$\left\langle n \right\rangle=m_{r}m_{l}A_{c}K_{a}$$

Here, $\boldsymbol{m}_{\mathbf{r}}$ is the CD40 site-density, $\boldsymbol{m}_{\mathbf{l}}$ is the αhCD40-SynNotch site density (converted from data in fig. S1B using calibration beads), and $\boldsymbol{A}_{\mathbf{c}}\boldsymbol{K}_{\mathbf{a}}$ is the effective 2D affinity determined in Fig. 2C. The above two equations are also listed on Fig. 2F.

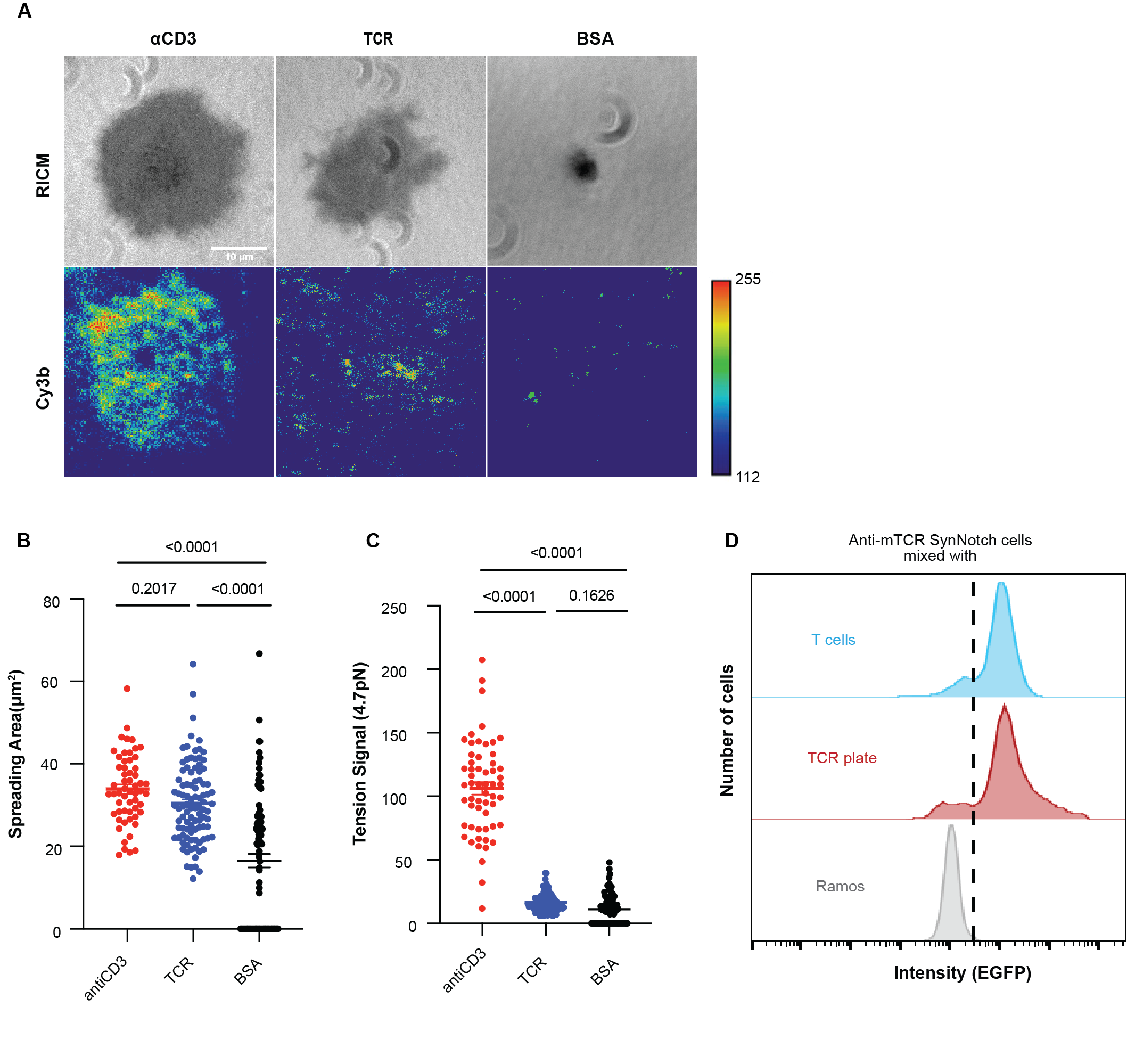

Fig. S4. The force on αmTCR-SynNotch was generated by the sender cell instead of the receiver cell. (A) Representative images by RICM and TIRF microscopy showing contact area (top row) and force signal (bottom row) of αmTCR-SynNotch expressing Jurkat cells 20 min after landing on coverslip surface functionalized by MTP tagged with anti-human CD3 (left column), mouse TCR (middle column), and BSA (right column). (B and C) Quantification of spreading area (B) and tension signal (C) exemplified in (A). Each point represents a cell, n = 57, 96, 92. (D) Histogram stack of EGFP fluorescence in αmTCR-SynNotch expressing Jurkat cells (3.4×10^4^) incubated either on a surface coated with 10 μg/ml soluble mouse TCR (middle) or with 1.66×10^5^ naïve CD8^+^ T cells (top, positive control) or with 1.66×10^5^ Ramos cells not expressing TCR (bottom, negative control) for 14 hours.

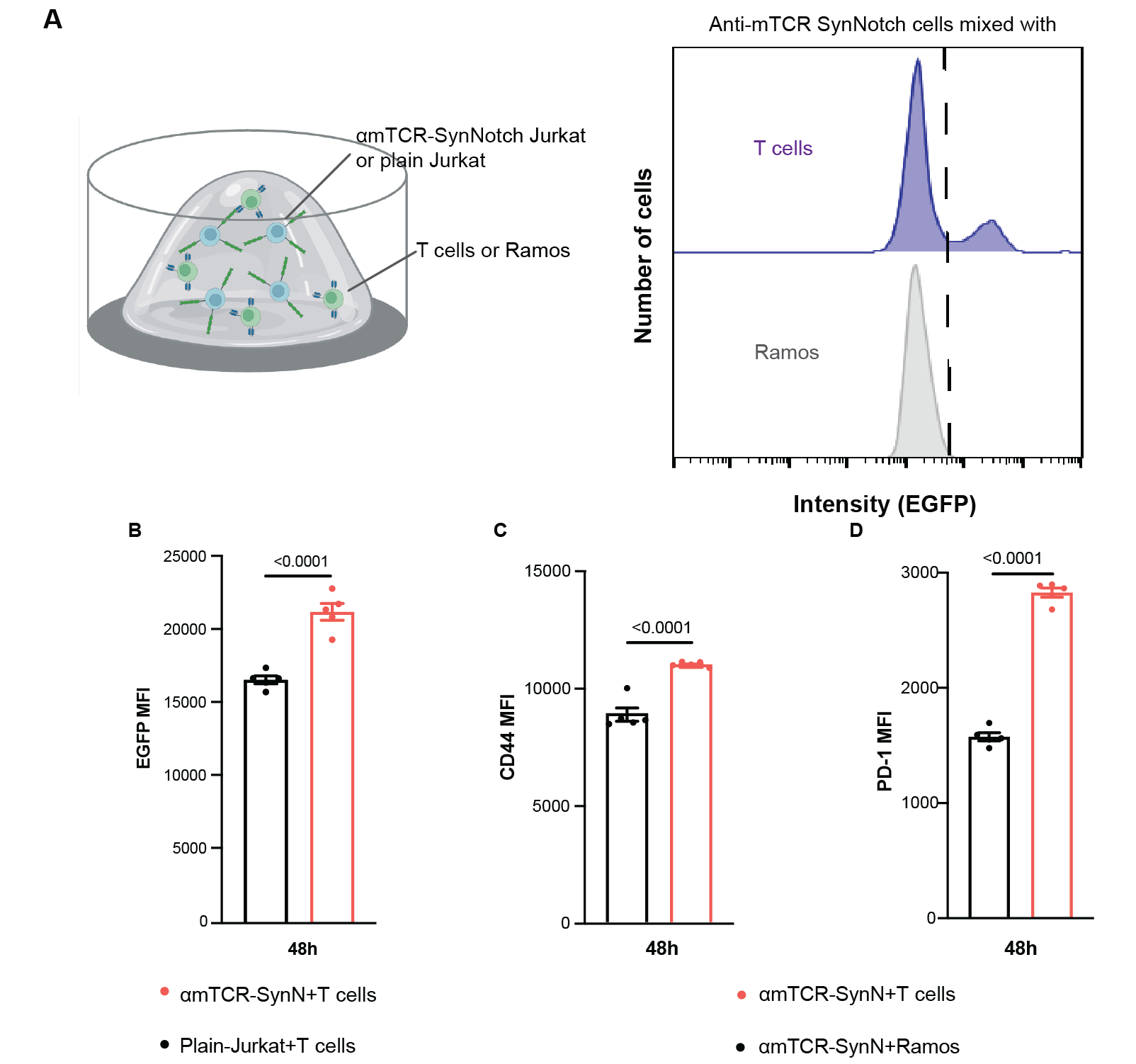

Fig. S5. Activation of αmTCR-SynNotch in 3D organoid coculture system. (A) Schematic of an organoid and the encapsulated cells (left) and fluorescence histograms showing results of the specificity control experiment (right). The experimental system and procedure that generated the results in (B to D) were similar to those in Fig. 4A except that smaller number of conditions were used, the sender cells and control cells were replaced by naïve CD8^+^ mouse T cells expressing mTCR and Romas B lymphoma cells not expressing mTCR, respectively, and the receiver cells (Jurkat) expressed αmTCR-SynNotch instead of αhCD40-SynNotch or not expressed SynNotch (plain Jurkat). 10^5^ αmTCR-SynNotch expressing Jurkat cells were encapsulated with 10^5^ naïve mouse T cells (blue) or 10^5^ negative control Ramos cells (gray) and cocultured in 3D organoids for 48 h. The organoids were degraded, the cells were harvested, and their EGFP expression level were measured using flow cytometry. (B) Mean ± SEM with individual data points (N = 5, n > 10,000) of EGFP MFI of αmTCR-SynNotch expressing Jurkat cells analyzed 48-h post coculture with mouse T cells (red) or Ramos cells (black). (C and D) Mean ± SEM with individual data points (N = 5, n > 10,000) of MFI of fluorescence staining of T cells by antibody against CD44 (C) or PD-1 (D) 48-h post coculture with Jurkat cells expressing (red) or not expressing (black) αmTCR-SynNotch. P-values were calculated using student-t tests.

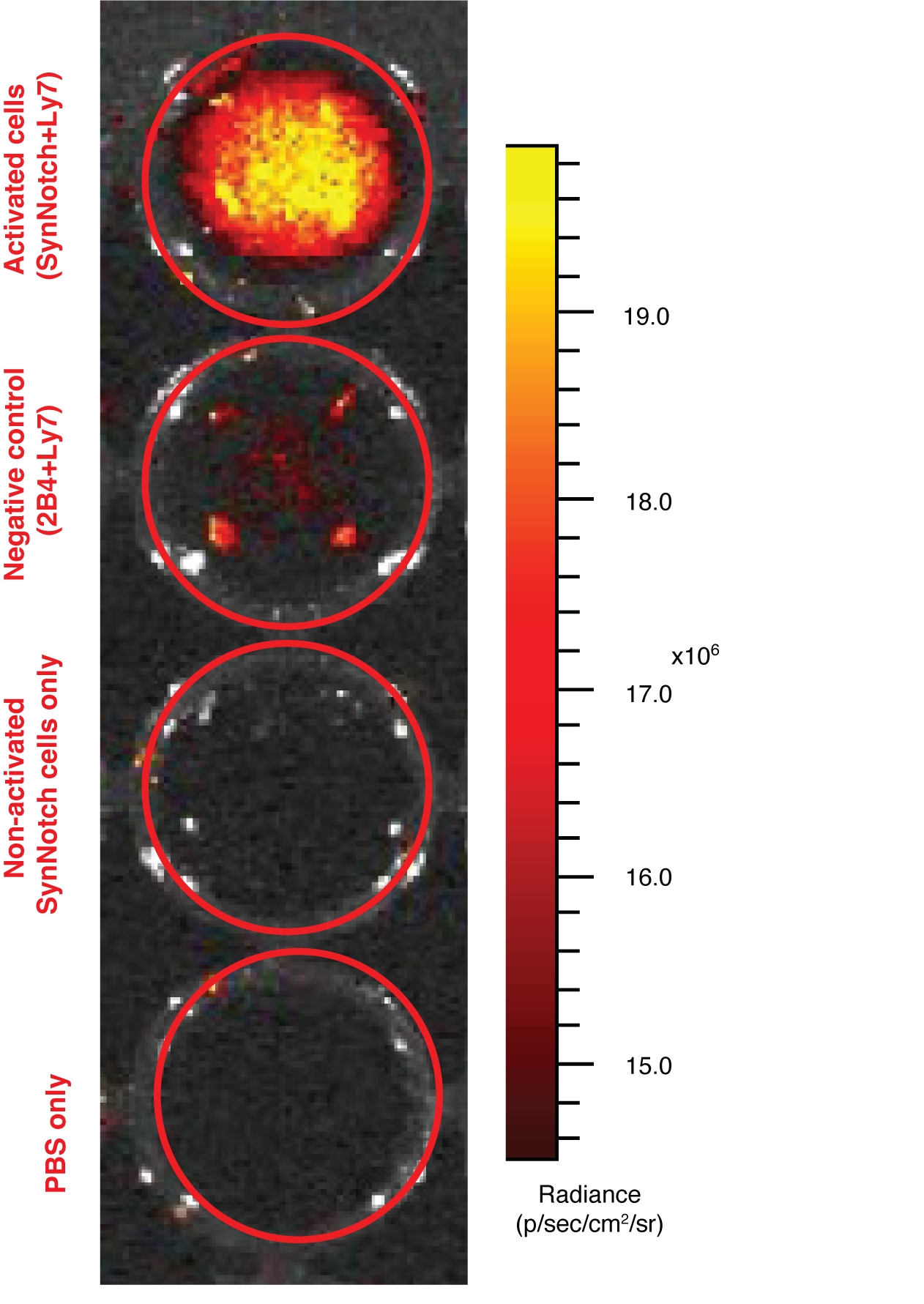

Fig. S6. Activated cells produce detectable fluorescence *in vitro* on IVIS. Cells (2×10^6^ each type) with indicated conditions were activated for ~16 h in a 1:1 ratio. Optimized Parameters for imaging: Exposure time: 5s| Binning factor: 4| Excitation: 500| Emission: 540| Fnumber: 1| FOV: 6.6.

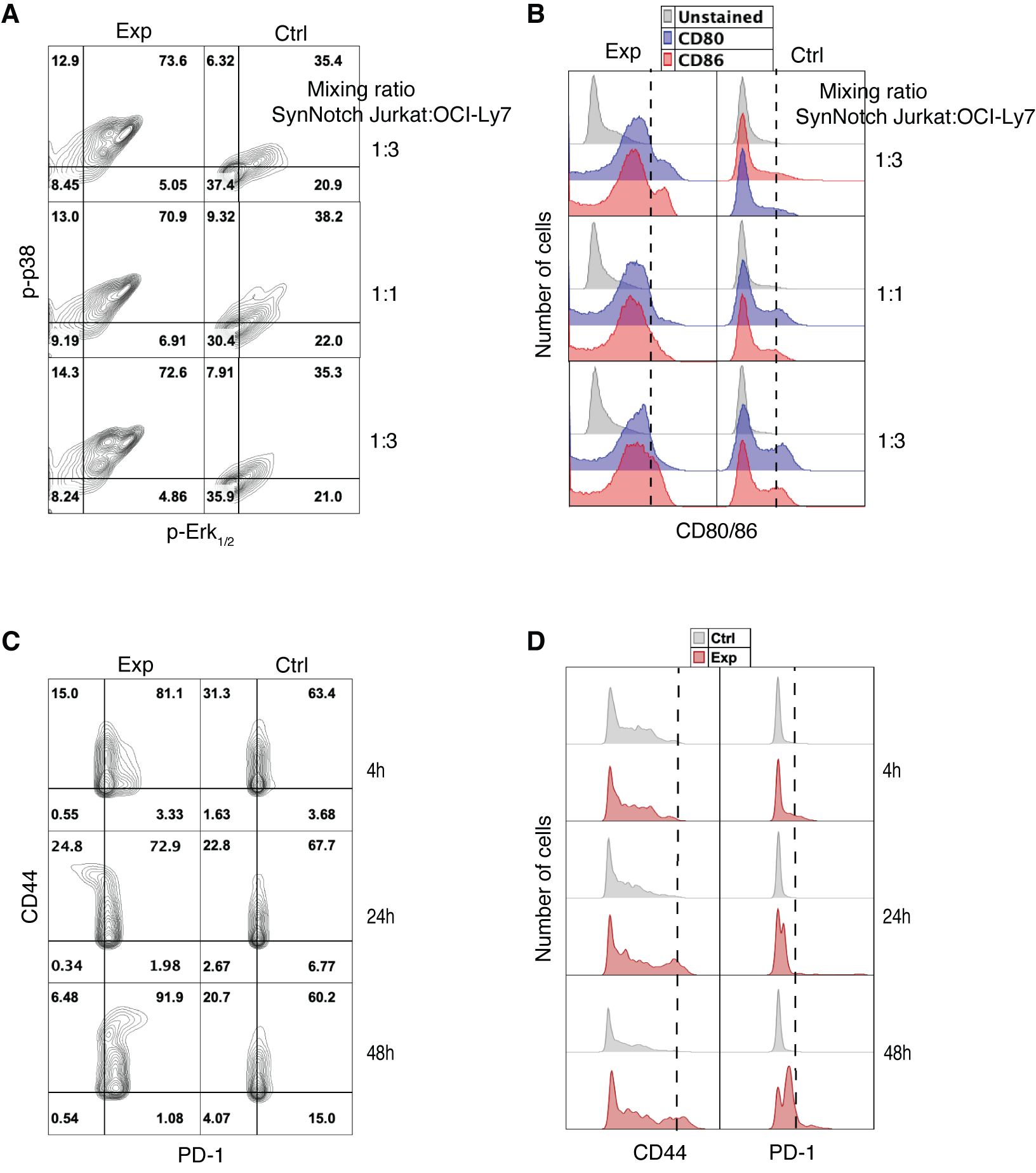

Fig. S7. In vivo activation of sender cells by receiver cells upon SynNotch engagement with target receptor. (A and B) Representative p-p38 vs p-Erk1/2 scatterplots (A) and CD80/CD86 histograms (B) of OCI-ly7 cells stained with fluorescently conjugated antibodies against p-p38 (y-axis in A), p-Erk1/2 (x-axis in A), CD80 (blue in B), CD86 (red in B), or unstained cells (gray in B) previously encapsulated with Jurkat cells expressing αhCD40-SynNotch (left column in B) or plain Jurkat cells (right column in B) at the indicated receiver/sender cell ratios. OCI-Ly7 cells were harvested and sorted from organoids previously implanted in NSG mice. (C and D) Representative CD44 vs PD-1 scatterplots (C) and CD44/PD-1 histograms (D) of mouse T cells stained with fluorescently conjugated antibodies against CD44 (y-axis in C and left column in D), PD-1 (x-axis in C and right column in D) previously encapsulated with Jurkat cells expressing αmTCR-SynNotch (red in D) or plain Jurkat cells (gray in D) at a 1:1 receiver/sender cell ratio. Mouse T cells were harvested and sorted from organoids previously implanted in NSG mice for the indicated times.

| REAGENT or RESOURCE | SOURCE | IDENTIFIER |
| --- | --- | --- |
| Antibodies |  |  |
| Myc-Tag (9B11) Mouse mAb (PE Conjugatge) | Cell Signaling Technology | 3739S |
| Myc-Tag (9B11) Mouse mAb (Alexa Fluor® 647 Conjugate) | Cell Signaling Technology | 2233S |
| PE anti-human CD40 antibody | Biolegend | 313005 |
| Biotin anti-human CD3 antibody, OKT3 | Biolegend | 317319 |
| Biotin anti-human CD40 antibody, 5C3 | Biolegend | 334343 |
| PE p44/42 MAPK (Erk1/2) (Thr202/Tyr204) antibody | Cell Signaling Technology | 14095 |
| PE anti- human CD19 antibody | Biolegend | 982402 |
| Phospho-p38 MAPK (Thr180/Tyr182) (3D7) Rabbit mAb | Cell Signaling Technology | 6908 |
| Mouse PD-1 Alexa Fluor 750-conjugated antibody | Bio-Techne | FAB77381S |
| APC Anti-Human/Mouse CD44 (IM7) | Cytek Biosciences | 20-0441 |
| PE anti-mouse CD8b Antibody | Biolegend | 126607 |
| Chemicals, peptides, and recombinant proteins |  |  |
| RPMI 1640 Medium | Corning | 10-040-CV |
| IMDM | Quality Biological | 112-035-101 |
| DMEM with L-Glutamine, 4.5g/L Glucose and Sodium Pyruvate | Corning | MT10013CV |
| Fetal Bovine Serum | R&D Systems | S11150H |
| Penicillin-Streptomycin | Thermo Fisher Scientific | 15070063 |
| HEPES | Thermo Fisher Scientific | 15630130 |
| Sodium Pyruvate | Thermo Fisher Scientific | 11360070 |
| MEM Non-Essential Amino Acids | Thermo Fisher Scientific | 11140050 |
| Biotinylated Bovine Serum Albumin | Thermo Fisher Scientific | 29130 |
| Phosphate Buffered Saline | Thermo Fisher Scientific | MT21040CM |
| Absolute Ethanol 200 Proof | Decon Labs | 2716 |
| 3-Aminopropyltriethoxysilane (APTES) | Thermo Fisher Scientific | AC430941000 |
| Hydrogen peroxide 30% | JT Baker | 2186-01 |
| Sulfuric Acid | MilliporeSigma | SX12445 |
| Sulfo-NHS-Acetate | Thermo Fisher Scientific | 26777 |
| mPEG-SC | Biochempeg | MF001023-2K |
| LA-PEG-SC | Biochempeg | HE039023-3.4K |
| TrypLE | Thermo Fisher Scientific | 12605028 |
| Poly-L-Lysine | Sigma | P4832 |
| HBSS (calcium, magnesium, no phenol red) | Thermo Fisher Scientific | 14025076 |
| Biotinylated Human CD40 / TNFRSF5 Protein, Avitag™,His Tag (MALS verified) | Acro Biosystems | CD0-H82E8-25ug |
| Azide-PEG-NHS Ester | Click Chemistry Tools | AZ103-100 |
| Maleimide-Streptavidin | Thermo Fisher Scientific | 21102 |
| MEM Non-Essential Amino Acids Solution | Thermo Fisher Scientific | 11140076 |
| Paraformaldehyde, 4% | Thermo Fisher Scientific | J19943-K2 |
| Streptavidin | Thermo Fisher Scientific | 434302 |
| Biotin-PEG (3500)-NHS Ester | CD Bioparticles | CDN1602 |
| Biotin | Sigma-Aldrich | B4501 |
| LB broth powder | RPI | L24060-500.0 |
| Lenti-X concentrator | Takara Bio | 631232 |
| Polybrene | Millipore Sigma | TR-1003-G |
| Custom 8.8nm Gold Nanoparticiles, Tannic Acid, 0.05mg/mL, 400ML | nanoComposix | NCX-Custom |
| Lipofectamine 3000 Transfection Reagent | Thermo Fisher | L3000008 |
| NEB Stable *E*. Coli | NEB | C3040 |
| EZ-link Sulfo-NHS-LC-Biotin | Thermo Fisher Scientific | 21327 |
| 4 arm Maleimide functionalized polyethylene glycol (PEG-4-MAL, 20kDa) | Layson Bio | N/A |
| VPM (GCRDVPM↓SMRGGDRCG peptide) | AAPPTec | N/A |
| Dithiothreitol (DTT) | Sigma Aldrich | DTT-RO |
| REDV oligopeptide (GREDVGC) | AAPPTec | N/A |
| Worthington Biochemical Corporation COLLAGENASE TYPE I | Fisher Scientific | NC9482366 |
| DAPT | MedChemExpress | HY-13027 |
| Histopaque^®^-1077 | Sigma-Aldrich | 10771 |
| Oligonucleotides |  |  |
| 4.7 pN hairpin strand  GTGAAATACCGCACAGATGCGTTTGTATAAATGTTTTTTTCATTTATACTTTAAGAGCGCCACGTAGCCCAGC | IDT | N/A |
| A21B strand /5AmMC6/ CGCATCTGTGCGGTATTTCACTTT/3Bio/ | IDT | N/A |
| BHQ2 strand /5ThioMC6-D/TTTGCTGGGCTACGTGGCGCTCTT/3BHQ_2/ | IDT | N/A |
| Cy3b strand  /5Cy3b/CGCATCTGTGCGGTATTTCACTTT/3Bio/ | Khalid Salaita lab | N/A |
| 15 mer locker strand (4.7 pN)  AAAAAACATTTATAC | IDT | N/A |
| Recombinant DNA |  |  |
| Plasmid: GA021-pHR_pGK_CD19scFv_ChimericNotch_G4VP64 | Wendell Lim lab | N/A |
| Plasmid: GA170-pHR-GAL4_GFP-PGK_BFP-HF_seq | Wendell Lim lab | N/A |
| Plasmid: GA170-pHR-GAL4_Luciferase-PGK_mcherry_seq | Gabriel Kwong lab | N/A |
| Commercial assay kits |  |  |
| The QuantiBRITE PE beads (PE-Quantitation Kit) | BD Bioscience | 340495 |
| LIVE/DEAD™ Fixable Blue Dead Cell Stain Kit | Thermo Fisher Scientific | L23105 |
| EasySep™ Human CD19 Positive Selection Kit II | Stemcell Technologies | 17854 |
| Software and algorithms |  |  |
| SnapGene | Dotmatics | https://www.snapgene.com/ |
| GraphPad Prism | GraphPad Software | http://www.graphpad.com/scientific-software/  prism/ |
| Flowjo | BD Bioscience | https://www.flowjo.com/ |
| Labview | National Instruments | http://www.ni.com/en-us.html |
| Matlab | MathWorks | https://www.mathworks.com/products/matlab.html |
| SolidWorks | Dassault Systems | https://www.solidworks.com/ |
| Other |  |  |
| Coverslip Mini-Rack, teflon | Thermo Fisher Scientific | C14784 |
| Attofluor Cell Chamber, for microscopy | Thermo Fisher Scientific | A7816 |
| Sticky-Slide VI 0.4 | Ibidi | 80608 |
| Dynabeads™ M-270 Streptavidin | Invitrogen | 65306 |
| Circle coverslips | Thermo Fisher Scientific | 64-0700 |
| Flow-Mix 5-Minute Epoxy | Devcon | 20455 |
| Non-culture tissued 24 well plates | Corning | 351147 |
| Sharp Tweezers | Thermo Fisher Scientific | 12-000-122 |
| Magnet | Supermagnetman | C0052 |
| Magnetic lid | In this paper | N/A |
| 96 Well Polypropylene Plates | Thermo Scientific | EK20201 |

Table S1. Resources used in this work. This table provides a complete list of all the reagents, assay kits, and software and algorithms used in this study, including antibodies, chemicals, peptides, recombinant proteins, oligonucleotides, recombinant DNA, critical commercial assays, software and algorithms.
